# MultiGS: A comprehensive and user-friendly genomic prediction platform Integrating statistical, machine learning, and deep learning models for breeders

**DOI:** 10.64898/2026.01.02.697306

**Authors:** Frank M. You, Chunfang Zheng, John Joseph Zagariah Daniel, Pingchuan Li, Bunyamin Tar’an, Sylvie Cloutier

## Abstract

Genomic selection (GS) is a core strategy in modern breeding programs, yet the rapid expansion of statistical, machine-learning (ML), and deep-learning (DL) models has made systematic evaluation and practical deployment increasingly challenging. To address these issues, we developed MultiGS, a unified and user-friendly framework that integrates linear, ML, DL, hybrid, and ensemble GS models within a standardized and computationally efficient workflow. MultiGS is implemented through two complementary pipelines: MultiGS-R, a Java/R pipeline implementing 12 statistical and ML models, and MultiGS-P, a Python pipeline integrating 17 models including five linear models, three ML approaches, and nine recently developed DL architectures implemented within the framework. We benchmarked MultiGS using wheat, maize, and flax datasets representing contrasting prediction scenarios. Wheat and maize were evaluated using random training–test splits within the same population, reflecting suitable conditions for assessing model capacity and scalability. Under these scenarios, several DL, hybrid, and ensemble models achieved prediction accuracies comparable to RR-BLUP and consistently exceeded those of GBLUP. In contrast, the flax dataset represented a true across-population prediction scenario with limited training set size and strong population structure. In this challenging context, classical linear models provided stable baselines, while a subset of DL architectures—particularly graph-based models and BLUP-integrated hybrids—demonstrated comparatively improved generalization across populations. Comparisons with previously published DL tools showed that MultiGS models achieved comparable or improved prediction accuracies while requiring lower computational costs, enabling routine retraining and large-scale evaluation. Overall, MultiGS informs, scenario-specific model selection and provides a practical platform for deploying genomic prediction under realistic breeding conditions. The software is freely available on GitHub (https://github.com/AAFC-ORDC-Crop-Bioinfomatics/MultiGS).

## INTRODUCTION

Genomic selection (GS) has become a core strategy in modern plant and animal breeding, enabling the prediction of breeding values using genome-wide markers and accelerating genetic gain through reduced cycle time (Meuwissen et al. 2001; Heffner et al. 2009). Since its introduction, a wide range of statistical, machine learning (ML), and deep learning (DL) models have been developed to improve prediction accuracy (PA) across diverse breeding scenarios. Traditional linear models such as Ridge Regression BLUP (RR-BLUP) and Genomic BLUP (GBLUP) remain widely adopted due to their simplicity, computational efficiency, and strong baseline performance (VanRaden 2008; Endelman 2011), while Bayesian approaches—including Bayesian Ridge Regression (BRR), Bayesian LASSO (BL), and BayesA/B/C—provide additional flexibility in modeling heterogeneous marker-effect distributions (de Los Campos et al. 2013; Perez and de los Campos 2014).

ML approaches such as Random Forest, Support Vector Machines, Gradient Boosting, and regularized regression have expanded the analytical landscape for GS (Gonzalez-Recio and Forni 2011; Heslot et al. 2012). More recently, DL architectures—including multilayer perceptrons (MLP), convolutional neural networks (CNN), recurrent networks, attention-based models, Transformers, and graph neural networks (GNN)—have shown promises for capturing nonlinear effects, epistasis, and structural dependencies among markers (Ma et al. 2018; Montesinos-López et al. 2018; Azodi et al. 2019; Sandhu et al. 2020). This led to a proliferation of DL-based GS methods, such as DeepGS (Ma et al. 2018), G2PDeep (Liu et al. 2019), DeepGP (Zingaretti et al. 2020), DNNGP (Wang et al. 2023), GPformer (Wu et al. 2023), Cropformer (Wang et al. 2025b), GEFormer (Yao et al. 2025), iADEP (Ye et al. 2025), WheatGP (Wang et al. 2025a), SoyDNGP (Gao et al. 2023) and DPCformer (Deng et al. 2025), each exploring different architectural choices and reporting improvements relative to traditional models.

Despite notable progresses, several limitations still hinder the broad adoption of DL–based GS methods in practical breeding pipelines. Many existing DL GS tools provide only partial or research-oriented implementations, with fragmented codebases, inconsistencies between published descriptions and actual source code, limited documentation, or incomplete end-to-end workflows. As a result, these tools often suffer from poor usability and limited reproducibility, making them difficult to integrate into real-world breeding programs. Many require substantial expertise in Python, PyTorch (Paszke et al. 2019), Graphic processing unit (GPU) computing, or Unix/Linux environments—skills that are uncommon among breeders and applied geneticists. As such, while DL models show strong potential, their accessibility to breeding programs remains limited.

A second challenge arises from inconsistent and often incomparable benchmarking practices across studies. Reported improvements over RR-BLUP, GBLUP, or third-party software frequently depend on inconsistent baseline implementations (e.g., R-based vs. Python-based models producing different accuracies), and disparate cross-validation strategies. Based on our experience while implementing both R and Python versions of standard GS models, even nominally identical baseline methods can produce large performance differences depending on software, preprocessing, and hyperparameters. Consequently, it is difficult to draw reliable conclusions about whether a DL model consistently outperforms existing approaches or merely performs well on a specific dataset under a particular configuration. The field lacks a unified, reproducible benchmarking platform that simultaneously supports diverse model families and standardized evaluation procedures.

To address these gaps, we developed MultiGS, a pair of complementary, user-friendly genomic prediction platform that 1) integrates a broad spectrum of GS models, ranging from classical linear mixed models to advanced deep learning (DL) and hybrid architectures in this study, 2) provide standardized workflows for data preprocessing, cross-validation (CV), across-population prediction (APP), and post-analysis, and 3) support SNP, haplotype, and principal component (PC) marker representations. In this paper, we describe the design and implementation of the MultiGS framework and evaluate its performance across multiple datasets. Our results show that the DL models implemented within the MultiGS framework consistently outperform GBLUP and achieve prediction accuracies (PAs) comparable to RR-BLUP, reinforcing recent evidence that DL approaches can match or exceed traditional GS models under appropriate conditions. Overall, MultiGS bridges methodological innovation and practical usability, offering an integrated platform to advance genomic selection research and facilitating integration of GS into practical breeding.

## MATERIALS AND METHODS

### Overview of the MultiGS framework

The MultiGS framework was developed to provide an integrated and reproducible platform for GS as the field expands from traditional mixed models towards increasing diverse ML and DL approaches. In particular, MultiGS places strong emphasis on the systematic evaluation of nine DL model archetectures that capture nonlinear, local, and graph-structured genotype–phenotype relationships beyond the assumptions of classical linear methods. Existing GS pipelines often require users to navigate multiple software tools with inconsistent workflows and incompatible preprocessing procedures, making fair comparison and practical deployment of DL models especially challenging. MultiGS resolves these limitations by offering a unified ecosystem that standardizes data handling, model execution, cross-validation, and result summarization across statistical, ML, and DL methodologies. The framework supports three marker representations derived from SNP genotypes: single-marker SNP representations (SNP), haplotype-based representations (HAP), and principal component (PC) representations (PC) and adopts a consistent input–output structure across all implemented algorithms, enabling direct benchmarking of classical models against advanced DL architectures.

MultiGS is organized into two complementary pipelines. MultiGS-R provides access to classical GS and Bayesian methods implemented in R serving as robust baselines widely used in breeding programs. MultiGS-P substantially extends the analytical scope by implementing advanced ML methods and nine DL models, including fully connected networks, convolution–attention hybrids, graph neural networks, and BLUP-integrated hybrid architectures. These DL models were designed to explicitly target key challenges in GS, such as nonlinear marker effects, local linkage disequilibrium (LD) patterns, and population structure, within a common and reproducible framework.

Both pipelines share the same design logic, allowing users to evaluate linear, ML, and DL models under identical preprocessing steps, training configurations, and accuracy metrics. Detailed descriptions of the R and Python pipelines are shown in Figure 1. To ensure user-friendliness, MultiGS employs a standardized input–output format controlled by a configuration file. Users can flexibly select any combination of models, marker representations, and evaluation modes (model benchmarking or prediction) simply by modifying configuration flags, without altering their data preparation workflow. This design enables breeders and researchers to focus on biological interpretation and breeding decisions while facilitating rigorous assessment of advanced DL models alongside established GS approaches.

**Figure 1.**
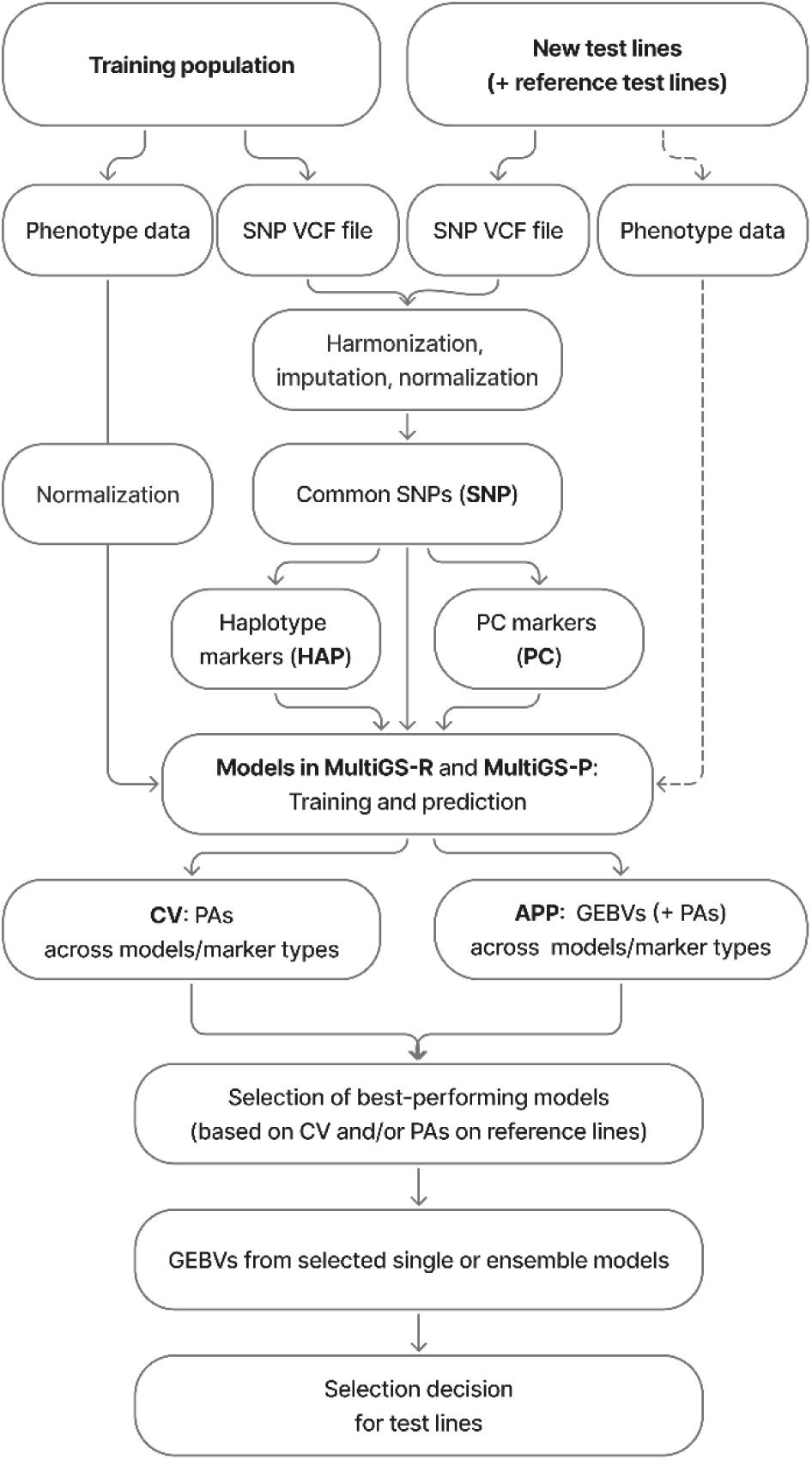
Schematic overview of the genomic prediction platform used to predict the genomic estimated breeding values (GEBVs) of new breeding lines with MultiGS-R and MultiGS-P. The reference test lines with phenotypic data and the dotted box containing phenotypic data from the test population are optional and used only for model evaluation and model selection when available. PA: prediction accuracy; CV: cross-validation; APP: across-population prediction; SNP: single nucleotide polymorphism; HAP: haplotype; PC: principal component; VCF: variant call format.

### MultiGS-R

MultiGS-R is the R-based pipeline for statistical and ML models. It integrates twelve widely used GS models through the *rrBLUP* (Endelman 2011), *BGLR* (Perez and de los Campos 2014), *e1071* (Meyer et al. 2023), and *randomForest* (Liaw and Wiener 2002) packages. These include linear mixed models such as RR-BLUP and GBLUP, Bayesian regression models including BRR, BL, and BayesA/B/C, random forest algorithms, support vector machines, and RKHS methods (Table S1). MultiGS-R automates the core steps of GS analysis—genotype and phenotype preprocessing, model training, prediction, and accuracy assessment—while maintaining consistent output formats across models. This pipeline provides a stable and accessible environment for breeders and researchers who require reliable and interpretable models without the need for extensive scripting.

### MultiGS-P

MultiGS-P implements an extensive suite of ML and DL models using Python, scikit-learn (Pedregosa et al. 2011), and PyTorch (Paszke et al. 2019). The pipeline includes linear and regularized models (RRBLUP-equivalent Ridge, Elastic Net, and Bayesian Ridge Regression), tree-based learners (Random Forest Regression), and gradient boosting methods (XGBoost and LightGBM) (Table S2). In addition to these conventional ML approaches, MultiGS-P implements nine recently developed DL architectures, ranging from fully connected networks and convolution–attention hybrids to multiple graph neural network variants (Table 1). The pipeline further integrates hybrid methods that combine RR-BLUP with deep networks (DeepResBLUP and DeepBLUP), as well as a stacking-based ensemble learner (EnsembleGS) capable of fusing predictions from arbitrary base models (Table 1). The detailed a Together, this collection provides a comprehensive modeling environment in which additive, nonlinear, LD-aware, graph-structured, and hybrid genotype–phenotype relationships can be evaluated within a unified and reproducible workflow.

**Table 1.** Summary of nine deep learning model architectures implemented in MultiGS-P

| Model | Architecture Type | Main Components | Key Properties |
| --- | --- | --- | --- |
| **DNNGS** | Fully connected deep neural network | Input dropout; 4 fully connected blocks (512–256–128–64) with ReLU and dropout; optional batch normalization; final linear prediction head | Efficient nonlinear modeling of genotype features; simple and fast baseline DL model |
| **MLPGS** | MLP with normalization and residual connections | Input dropout; 2 dense blocks (128→64) with GELU/ReLU, dropout, layer normalization; optional residual skip connections; output normalization + final dense layer | Stabilized deep MLP with improved training dynamics and gradient flow |
| **GraphConvGS** | Graph Convolutional Network (GCN) | KNN graph construction; 2×GCNConv + layer norm + ReLU + dropout; node-wise MLP | Models sample-to-sample similarity; effective for population-structured GS data |
| **GraphAttnGS** | Graph Attention Network (GAT) | KNN graph; 2×GATConv (multi-head attention) + layer norm + dropout; node MLP | Learns adaptive attention weights over neighbors; flexible modeling of heterogeneous relationships |
| **GraphSAGEGS** | GraphSAGE neighborhood aggregator | KNN graph; 2×SAGEConv + layer norm + dropout; node MLP | Inductive, scalable, and robust across populations; robust performance on large datasets |
| **GraphFormer** | GraphSAGE + Transformer hybrid | 2×SAGEConv → node embeddings → Transformer encoder → MLP | Captures both local graph structure and global node interactions for enhanced representation learning |
| **DeepResBLUP** | Residual hybrid (RRBLUP + DL) | Fit RRBLUP baseline; DL model fits residual signal; weighted combination of linear and nonlinear predictions | Effective additive baseline with nonlinear correction; interpretable and stable |
| **DeepBLUP** | Integrated BLUP-in-DL architecture | RRBLUP-like linear layer → 3 dense blocks (256–128–64) with GELU, batch norm, dropout, and residual connections; optional skip link from RRBLUP output | Deep refinement of BLUP with modern DL structure; robust performance for additive + mild nonlinear effects |
| **EnsembleGS** | Stacked ensemble | Trains multiple base models; collects OOF predictions; meta-learner (linear regression) for final prediction | Most robust across datasets; leverages complementary strengths of diverse model families |

BLUP, best linear unbiased prediction; BN, batch normalization; Conv, convolution (layer); DL, deep learning; FC, fully connected (layer); FFN, feed-forward network; GCN, graph convolutional network; GAT, graph attention network; GELU, Gaussian Error Linear Unit; LD, linkage disequilibrium; KNN, k-nearest neighbors; MHA, multi-head attention; MLP, multilayer perceptron; OOF, out-of-fold; ReLU, rectified linear unit; SAGE, sample and aggregate (GraphSAGE)

All DL models were systematically tuned during development to ensure stable training and competitive performance across datasets. The default hyperparameters for all models implemented in MultiGS-P are provided in the template configuration files and are summarized in **Table S3**.

The DL architectures are grouped into fully connected, graph-based, hybrid, and BLUP-integrated categories. Each architecture is described in the Supplementary Materials and Methods with documentation of its design rationale and intended uses (**Figures S1** and **S2**). All ML and DL models in MultiGS-P are fully configurable through a centralized configuration file (**Table S3**), allowing users to adjust hyperparameters, model depth, learning schedules, and regularization settings without modifying source code. This design facilitates systematic benchmarking and fair comparison across diverse model classes while supporting flexible adaptation to a range of datasets and breeding scenarios.

### Hyperparameter tuning of deep-learning models

All deep learning (DL) models were tuned during development using the flax training dataset, which represents a challenging scenario with a small training population and strong population structure and therefore better reflects realistic breeding conditions than within-population validation. Hyperparameters identified under this conservative setting were fixed and applied uniformly to all datasets (wheat2000, maize6000, and flax287). No dataset- or trait-specific tuning was performed, ensuring fair and consistent comparison among model classes.

Hyperparameter tuning was conducted using a grid search strategy, and the resulting default settings are provided in the template configuration files and summarized in Supplementary Table S1. For users who wish to optimize models for specific datasets, a utility script (hyperparamer_optimizer.py) is provided to facilitate targeted hyperparameter optimization of major model parameters

### Marker representations

MultiGS supports three genomic feature representations—SNP, HAP, and PC—allowing users to evaluate model performance across alternative marker representations. These representations capture complementary aspects of genomic variation and provide flexibility for modeling different trait architectures.

SNP representation uses the raw genotype matrix, encoding markers as additive allele counts. Although referred to here as SNP-based, this representation can accommodate any bi-allelic or multi-allelic markers derived from different genotyping technologies, provided they are numerically encoded. This representation is the most widely used in genomic selection and serves as the baseline input for most models.

Haplotype representation aggregates adjacent SNPs into haplotype blocks, enabling models to capture local linkage-phase information and multi-allelic patterns that may correlate more strongly with causal loci. This view is particularly useful in regions with strong LD or when phased or block-based genotype data are available. Both pipelines used additive dosage coding with values 0, 1, and 2: code 0 (0/0) for homozygous reference alleles, code 1 (0/1, 0/2, …) for heterozygous genotypes with one alternative allele, and code 2 (1/1, 2/2, …) for homozygous alternative alleles. Missing data (./.) are assigned code –1, and are imputed using the mean algorithm or Beagle software (Browning et al. 2021). Haplotype blocks were estimated with the *rtm-gwas-snpldb* tool in RTM-GWAS v2020.0 (He et al. 2017), which applies the Haploview “Gabriel” confidence-interval algorithm (Gabriel et al. 2002; Barrett et al. 2005). This algorithm identifies haplotype blocks based on statistically supported LD confidence intervals and has been widely adopted for defining robust LD blocks. Pairwise LD was measured using D’ with 95% confidence intervals, and haplotype blocks were defined when ≥95% of informative pairs were in strong LD (CI(D’) lower ≥ 0.70, upper ≥ 0.98).

PC representation summarizes genome-wide marker information using PCs derived from SNP data. PCs capture major population structure and relatedness patterns while reducing dimensionality. This view provides a compact genomic representation and may improve stability or computational efficiency in certain models. The first *N* PCs explaining 95% (configurable) of the total variance is retained for prediction. The number of retained PCs varied across datasets but typically ranged between ∼20 and 200 PC.

All three marker representations are automatically constructed within both MultiGS pipelines, ensuring consistent preprocessing and compatibility across model families. Marker views are stored using a unified format, enabling direct comparison of PAs across different genomic representations.

### Model evaluation

MultiGS provides standardized evaluation procedures to ensure reliable comparison of models, marker representations, and data partitions. Both pipelines implement CV schemes commonly used in genomic prediction studies. These include random k-fold CV for assessing general predictive performance within populations and structured CV options suitable for breeding programs involving multiple families or subpopulations. CV routines are fully automated, with users specifying fold number and seed settings to maintain reproducibility. In addition to within-population evaluation, MultiGS supports across-population prediction, a scenario in breeding applications where training and target populations differ.

Pearson’s correlation coefficient between predicted and observed values, hereafter referred to as prediction accuracy (PA), was used consistently as the performance metric across all models. MultiGS also generates prediction summaries and visualizations that facilitate rapid comparison of model performance. Result objects are standardized across both pipelines, allowing seamless benchmarking of traditional statistical models against ML and DL approaches.

### Case studies

To evaluate the performance of the models implemented in the MultiGS pipelines, three genomic and phenotypic datasets—wheat, maize, and flax—were used. The wheat dataset consisted of 2,403 Iranian bread wheat (*Triticum aestivum*) landrace accessions conserved in the CIMMYT Wheat Gene Bank (https://hdl.handle.net/11529/10548918) (Crossa et al. 2016). Genotyping was performed using 33,709 DArTseq presence/absence markers, coded as 1 (allele present) or 0 (allele absent). Five traits—thousand-kernel weight (TKW), grain width (GW), grain hardness (GH), grain protein (GP), and grain length (GL)—were evaluated. After removing accessions with missing phenotypes, the remaining 2000 accessions were randomly divided into a training population of 1,600 and a testing population of 400 (80:20 ratio). After filtering using minor allele frequency ≥ 5%; call rate ≥ 80%, the retained 9,927 markers were converted to VCF format for genomic prediction analyses. This dataset is hereafter referred to as wheat2000.

The maize dataset was derived from the CUBIC hybrid population described by Liu et al. (2020) . In total, 6,210 F₁ hybrids (207 female × 30 male) were evaluated across five environments in China during the 2015 growing season. Three agronomic traits—days to tassel (DTT), plant height (PH), and ear weight (EW)—were measured, and BLUPs were calculated within the 2015 trials. Genomic data consisted of 10,000 SNPs randomly sampled from ∼4.5 million imputed markers available through the ZEAMAP repository (https://ftp2.cngb.org/pub/CNSA/data3/CNP0001565/zeamap/99_MaizegoResources/01_CUBI <u>C_related/</u>). After removing hybrids missing genotypic or phenotypic records, the dataset comprised 5,831 maize hybrids. These hybrids were randomly partitioned into a training population of 4,664 hybrids (80%) and a testing population of 1,167 hybrids (20%). For convenience, this dataset is hereafter referred to as maize6000.

To assess the capacity of the MultiGS models to perform prediction across populations (as opposed to cross-validation), we used a flax dataset comprising 278 linseed accessions that had been sequenced using a whole-genome shotgun approach (hereafter referred to as flax287) (You et al. 2018; He et al. 2023; You et al. 2017), yielding approximately 1.7 million SNPs for model training. To mimic a practical breeding scenario relevant to variety development, a test population consisting of 260 inbred lines derived from multiple biparental populations (You et al. 2018) was used. These test lines were re-genotyped by mapping short pair-ended reads to the latest flax reference genome, CDC Bethune v3.0 (NCBI accession JAZBJT000000000; You et al., in preparation), yielding 43,179 SNPs, of which 33,895 were shared with the training populations and used for genomic prediction. Three key agronomic traits—days to maturity (DTM), oil content (OIL), and plant height (PH)—were evaluated to assess predictive performance.

The genetic diversity and population differentiation between training and test sets across these three datasets are summarized in Table S4. Wheat2000 and Maize600 exhibited negligible genetic differentiation between training and test populations (FST ≈ 0), whereas Flax287, representing a true across-population prediction scenario, showed strong population divergence (FST = 0.27). In addition, expected heterozygosity (He) was high in maize6000 (∼0.38) but markedly lower in wheat2000 and flax287 (∼0.01-0.02), reflecting substantial differences in underlying population genetic structure among the datasets.

To benchmark the performance of the MultiGS framework against existing DL–based GS approaches, we selected four publicly available GS tools representing diverse neural-network architectures: DeepGS (convolutional neural networks) (Ma et al. 2018), CropFormer (CNN integrated with self-attention) (Wang et al. 2025b), WheatGP (CNN-LSTM–based feature extractors) (Wang et al. 2025a), and DPCFormer (Deng et al. 2025). Because these tools differ in their required input formats, preprocessing workflows, and output interfaces, direct comparison is not straightforward. To ensure consistent evaluation across models, we developed customized wrapper programs for each tool. These wrappers harmonized SNP marker data and phenotypic inputs, including: (i) aligning samples between genotype and phenotype files, (ii) identifying common markers between training and testing sets, (iii) converting VCF, comma separated value (CSV), or matrix formats into tool-specific input structures, and (iv) standardizing prediction outputs for downstream comparisons. All tools were run using identical training/test splits and the same set of single-trait prediction tasks. This unified preprocessing and execution framework ensures a fair, reproducible comparison between MultiGS models and third-party GS methods regardless of their heterogeneous interfaces.

To ensure fair comparison with previously published DL tools that require fixed input dimensions, dataset-specific adaptations were applied where necessary. For WheatGP, the original implementation hard-coded the Long Short-Term Memory (LSTM) input size as *lstm_dim* = 10,080, corresponding to the reference dataset used in the original study. However, the effective dimensionality of the LSTM layer depends on the number of markers provided as input. To enable application across datasets with varying SNP counts, we modified the implementation by replacing the fixed input size with a dynamic formulation, 𝑙𝑠𝑡𝑚_ dim = 8(𝑔_𝑖_ − 4), where 𝑔_𝑖_ denotes the number of SNP markers in the *i-th* group (five groups in total, as defined in WheatGP). This modification allows the LSTM input dimension to scale automatically with the number of available markers, enabling consistent application across datasets without altering model structure.

For CropFormer, the number of input markers is fixed at 10, 000 by design. To satisfy this constraint, random SNP subsetting or zero-padding was applied as appropriate. Specifically, when datasets contained more than 10,000 markers, a random subset of 10,000 SNPs was sampled; when fewer markers were available, padding was used to reach the required input size. The same procedure was applied consistently across the wheat2000, maize6000, and flax287 datasets. Random sampling was performed using fixed random seeds to ensure reproducibility.

## RESULTS

### Prediction accuracy (PA) across models in the wheat2000 dataset

Using the wheat2000 dataset (1,600 training and 400 independent test lines), we evaluated all models implemented in MultiGS-P and MultiGS-R and compared their performance with several published DL methods (**Figure 2**; **Table S5**). Linear mixed-model baselines (RR-BLUP, GBLUP, BRR) provided consistent reference performance across pipelines, with SNP-based PAs ranging from ∼0.53 to 0.76 depending on trait. Nearly identical results across implementations confirmed reproducibility, while BGLR-based GBLUP slightly outperformed rrBLUP-based implementations, consistent with known differences in variance component estimation.

**Figure 2.**
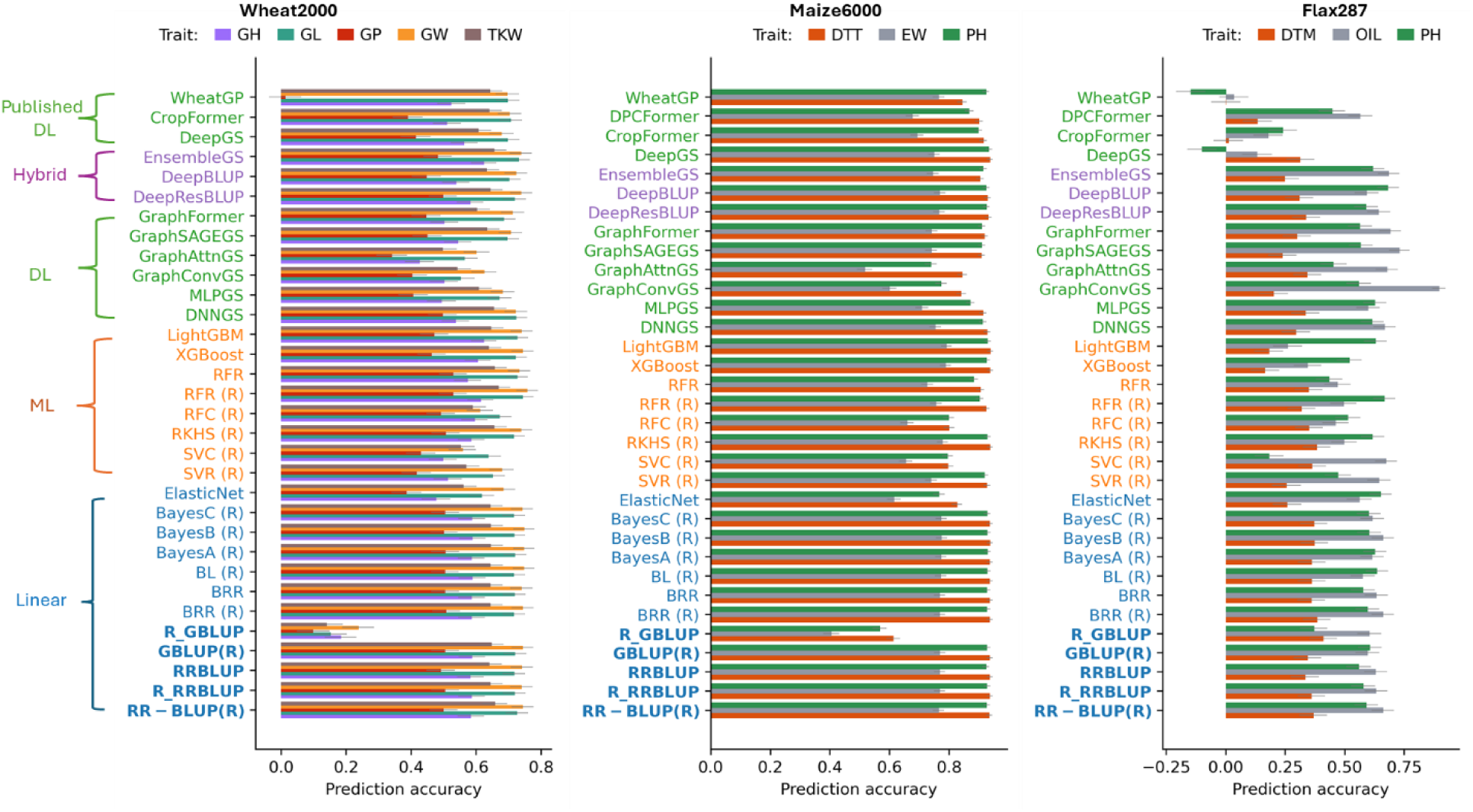
Prediction accuracies (PAs) across 17 models implemented in MultiGS-P, 12 models implemented in MultiGS-R (indicated by “(R)”), and several previously published third-party deep learning models for three datasets. Standard errors of PA are indicated on each bar. Models are categorized into Linear (blue), Machine Learning (orange), Deep Learning (green), and Hybrid (purple). RR-BLUP and GBLUP are highlighted in bold as baseline models. GH, grain hardness; GL, grain length; GP, grain protein; GW, grain width; TKW, thousand-kernel weight; DTT, days to tassel; EW, ear weight; PH, plant height; DTM, days to maturity; OIL, oil content; PH, plant height. Baseline models are shown in bold.

Several ML models matched or exceeded linear baselines, particularly tree-based approaches. Random Forest, XGBoost, and LightGBM frequently ranked among the strongest performers, achieving PAs of ∼0.72–0.74 for GL and GW, comparable to or slightly exceeding RR-BLUP. ElasticNet showed stable but generally weaker performance and did not consistently outperform linear models.

DL models showed competitive but trait-dependent performance. For high-heritability traits (GL, GW, TKW; h² ≈ 0.83–0.88), most DL architectures achieved PAs comparable to top ML models and RR-BLUP (typically ∼0.70–0.74). Among these, GraphSAGEGS and GraphFormer consistently outperformed other graph-based models, while DNNGS showed stable performance across traits. For lower-heritability traits (GH, GP), DL models slightly underperformed linear baselines but still achieved reasonable accuracies (∼0.45–0.60).

Hybrid approaches integrating RR-BLUP with DL (DeepBLUP, DeepResBLUP) ranked among the strongest DL-based models, with DeepResBLUP matching or slightly exceeding RR-BLUP for GL, GW, and TKW. The stacking-based EnsembleGS achieved consistently high accuracy across all traits (up to ∼0.74), highlighting the benefit of combining complementary predictors.

Compared with published DL tools (DeepGS, CropFormer, WheatGP), the top MultiGS models achieved comparable or higher PAs across all traits. EnsembleGS, DeepResBLUP, and GraphSAGEGS consistently matched or outperformed DeepGS, while CropFormer and WheatGP showed more variable performance, particularly for GP.

### Prediction accuracy (PA) across models in the maize6000 dataset

With the maize6000 dataset (4,664 training and 1,167 test lines), we evaluated 17 MultiGS-P and 12 MultiGS-R models using SNP-, HAP-, and PC-based markers across three traits (DTT, EW, PH) (**Figure 2**; **Table S6**). Linear baselines again performed strongly, particularly for DTT and PH, with SNP- and haplotype-based PAs consistently exceeding 0.92. For EW, accuracies were lower (∼0.76–0.77) but remained among the strongest results. Across traits, haplotype markers consistently outperformed PCs, reflecting the importance of local LD in maize.

Tree-based ML models performed particularly well. XGBoost and LightGBM achieved accuracies up to ∼0.94 for DTT and ∼0.93 for PH, matching or slightly exceeding RR-BLUP, and ∼0.78–0.79 for EW. In contrast, classification-oriented models (SVC, RFC) consistently underperformed for these continuous traits.

DL models achieved accuracies comparable to linear baselines for DTT and PH (typically ∼0.91–0.93). GraphSAGEGS, GraphFormer, DeepResBLUP, and DeepBLUP consistently ranked among the strongest DL models. For EW, performance was more variable; however, hybrid models (DeepResBLUP, DeepBLUP) matched RR-BLUP (∼0.76–0.77), and GraphSAGEGS and EnsembleGS also performed well.

Relative to published DL tools (DeepGS, CropFormer, DPCFormer, WheatGP), the best MultiGS models achieved comparable or improved accuracies across all traits. EnsembleGS, DeepResBLUP, DeepBLUP, and GraphFormer matched or slightly exceeded DeepGS and CropFormer for DTT and PH, while DeepBLUP and DeepResBLUP performed comparably to the strongest third-party models for EW. DPCFormer showed lower and more variable accuracy across traits. Overall, MultiGS achieved competitive DL performance while supporting a broader and more flexible model set.

### Prediction accuracy (PA) of models for across-population prediction in the flax287 dataset

The flax287 dataset represents a substantially more challenging scenario, combining a small training population (278 accessions) with prediction in a genetically narrow biparental population (260 lines) exhibiting strong population structure and genetic differentiation between training and test lines (**Figure S3, Table S4**). Across traits (DTM, OIL, PH), prediction accuracies were lower and more variable than in wheat2000 and maize6000 (**Figure 2**; **Table S7**). Haplotype-based models consistently outperformed SNP and PC encodings, reflecting strong LD structure in flax.

For DTM, all models showed limited predictive ability (HAP-based PAs ∼0.20–0.41). Linear baselines were among the most stable performers (∼0.33–0.41). ML models did not improve upon these results, and DL models showed high variability, with some architectures achieving baseline-level accuracy while others performed poorly or negatively. Published DL models exhibited similarly weak performance.

In contrast, th PAs for OIL were higher cross all model families. Linear models achieved PAs of ∼0.60–0.66, while several ML and DL models exceeded these baselines. DNNGS, GraphSAGEGS, GraphFormer, and EnsembleGS achieved accuracies up to ∼0.70–0.75, and GraphConvGS reached the highest overall PA (∼0.90). Hybrid models also performed well across marker representations. Among third-party DL tools, only DPCFormer showed moderate performance, while others transferred poorly to this across-population setting.

For PH, baseline accuracies reached ∼0.61, with several DL and hybrid models exceeding this level. DNNGS, GraphFormer, and DeepBLUP achieved PAs up to ∼0.66–0.70, while ElasticNet and LightGBM also performed competitively. Third-party DL models again showed inconsistent or weak performance.

Overall, the flax results underscore the strong effects of training population size, population structure, and marker representation on genomic prediction. Linear models provided stable baselines, while selected DL and hybrid models—particularly those incorporating additive genetic priors or graph-based representations—offered clear advantages for OIL and PH. These findings emphasize that DL benefits are trait- and context-dependent and support the value of a diverse modeling portfolio within the MultiGS framework.

### Effect of marker representation on prediction accuracy (PA)

For the three datasets, marker representation had a consistent and measurable impact on PAs across the 30 models implemented in MultiGS (**Figure 3**). For the wheat2000 dataset, where haplotype construction was not possible due to the absence of chromosome-level SNP coordinates, PAs based on SNPs and PCs were largely comparable, with only marginal differences regardless of the traits (average PA: SNP = 0.597, PC = 0.586). Trait-specific differences were similarly small), and SNP- and PC-based predictions differed by only 0.01–0.02 for the five traits, confirming that both marker representations captured similar predictive signals in this dataset.

**Figure 3.**
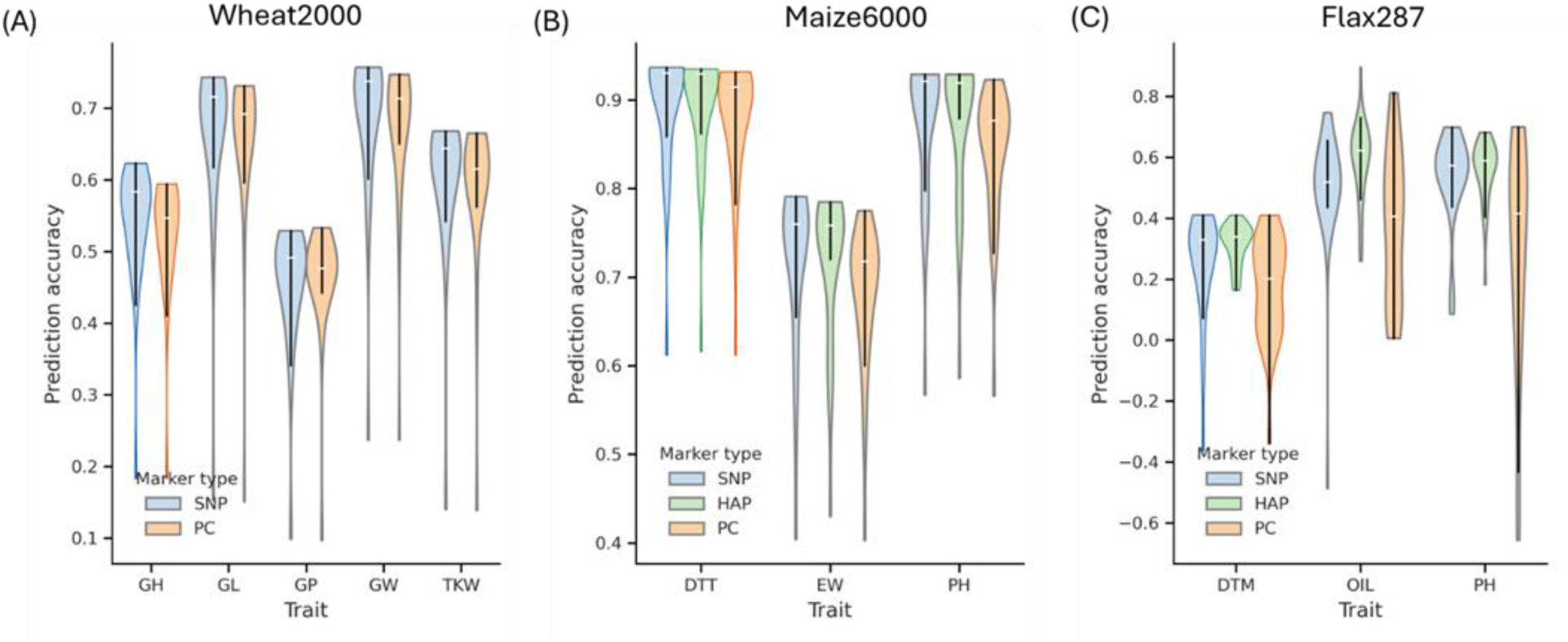
Prediction accuracies of three marker representations—single nucleotide polymorphisms (SNP), haplotypes (HAP), and principal components (PC)—across the 29 models implemented in both MultiGS pipelines in different datasets. **(A)** the wheat2000 dataset, **(B)** the maize6000 dataset, and **(C)** the flax287 dataset. GH, grain hardness; GL, grain length; GP, grain protein; GW, grain width; TKW, thousand-kernel weight; DTT, days to tassel; EW, ear weight; PH, plant height; DTM, days to maturity; OIL, oil content.

In contrast, for the maize6000 and flax287 datasets, where haplotype markers could be derived, HAP markers consistently outperformed both SNP and PC markers. In maize, haplotypes provided the highest accuracies across all three traits assessed (average PA: HAP = 0.831 vs. PC = 0.800), with notable improvements in EW (0.719 vs. 0.680) and PH (0.878 vs. 0.842). A similar trend was observed in flax, where HAP markers showed higher accuracies for DTM, OIL, and PH, improving PA by 0.12–0.26 over SNPs and by 0.24–0.28 over PCs. SNP markers ranked second in both datasets, while PC-based predictions were consistently the lowest, showing greater variability across traits and models.

Haplotype markers outperform or match SNP markers regardless of the dataset or traits, highlighting their value in capturing local LD and multi-allelic variations. PC markers showed dataset-dependent behavior, performing comparably to SNPs in wheat but underperformed compared to HAP and SNP markers in maize and flax. While PCs can provide a compressed representation of genomic structure, their use comes at sacrifice of predictive resolution that can be achieved with richer marker encodings such as haplotypes are available.

### Effect of Training Population Size on Prediction Accuracy

The relationship between PA and training population size was evaluated using the maize6000 dataset, which provides training sets ranging from 500 to 4,500 individuals. Seven representative models were assessed, including linear (R_RRBLUP), ML (LightGBM), DL (MLPGS, DNNGS, GraphSAGEGS), and hybrid approaches (DeepBLUP, EnsembleGS). Across all three traits (DTT, EW, and PH), PA increased consistently with training population size, with rapid gains observed between ∼500 and 2,500 samples and continued improvement at larger sizes (**Figure 4**).

**Figure 4.**
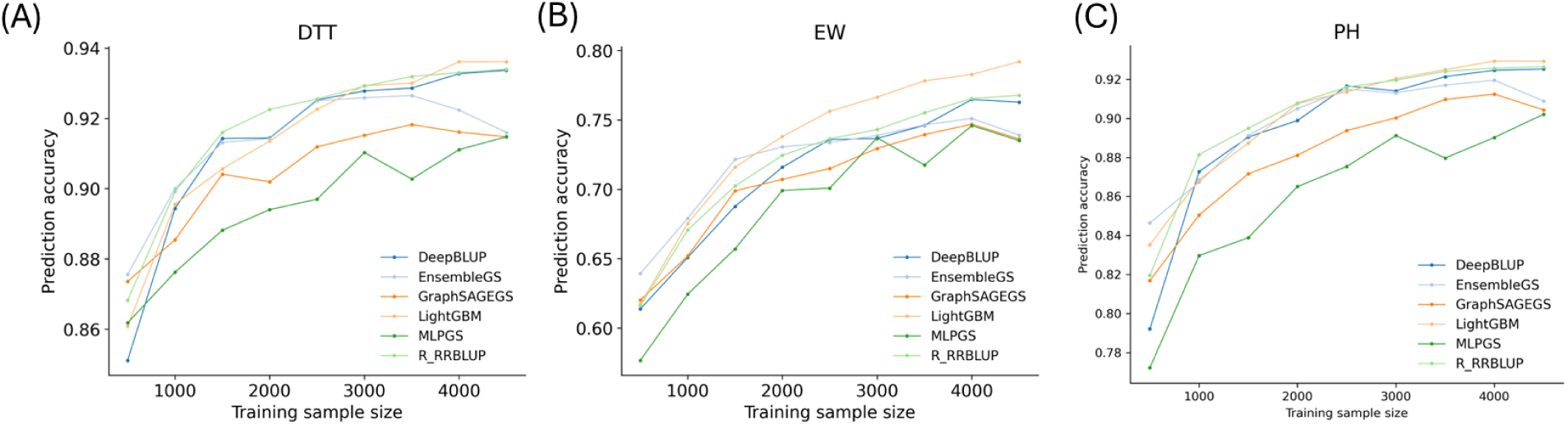
Prediction accuracies of three traits in the maize6000 dataset across varying training sample sizes. The full dataset contained 5831 samples, from which 4,664 lines (80%) were randomly selected as the training population and 1167 lines (20%) as the test set. A random subset of 10,000 SNPs was used as markers. From the 4,664 training lines, subsets of different sizes were randomly sampled and used to predict the fixed set of 1,167 test samples. Panels show results for: **(A)** days to tassel (DTT); **(B)** ear weight (EW); and **(C)** plant height (PH).

Although deep learning models are often assumed to require large datasets, linear models also benefited from increased sample size. R_RRBLUP achieved its highest accuracies at the largest training sizes and, in several cases, matched or exceeded DL models beyond 4,000 samples. LightGBM showed strong scalability and frequently performed best for traits with pronounced nonlinear components, particularly EW. The hybrid model DeepBLUP exhibited stable gains comparable to R_RRBLUP, whereas other DL models generally underperformed relative to the linear baseline across most training sizes. Overall, these results demonstrate that increased training population size leads to continued improvements in genomic prediction accuracy across all model classes, regardless of model complexity.

### Runtime performance across models

Using the maize6000 dataset, we compared the computational efficiency of 17 models implemented in MultiGS-P across linear, machine-learning (ML), deep-learning (DL), and hybrid categories under both CPU and GPU environments (**Figure 5A,B**), and evaluated four previously published DL models separately (**Figure 5C**). Runtime represents the average training time across three traits.

**Figure 5.**
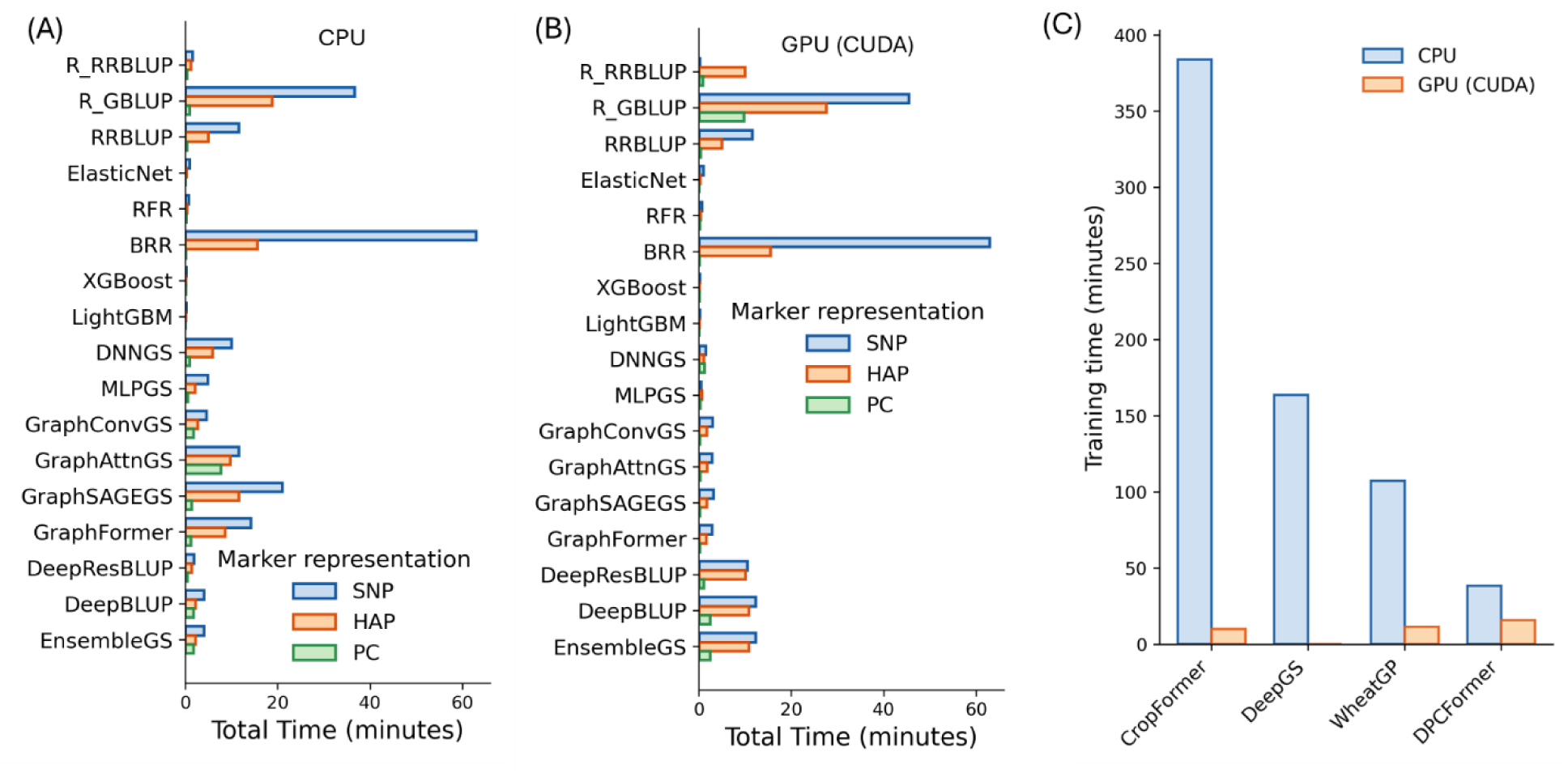
Runtime performance of deep learning models compared with linear and machine-learning models under CPU and GPU environments on the high-performance computing server. A total of 17 models implemented in MultiGS-P and four third-party tools were evaluated using the maize6000 dataset. Runtime represents the average training time required for model fitting of each trait. All benchmarks were conducted using the maize6000 dataset across three traits. **(A) and (B)** Runtime (minutes) for 17 models under CPU and GPU environments, respectively. **(C)** Runtime (minutes) for four previously published third-party deep learning models using SNPs.

Under CPU-only conditions, most linear and ML models completed training within seconds to a few minutes across marker types, with RR-BLUP, ElasticNet, Random Forest, XGBoost, and LightGBM showing consistently short runtimes. In contrast, Bayesian linear models (BRR and R_GBLUP) required substantially longer runtimes, in some cases exceeding those of several DL models, reflecting the computational cost of iterative Bayesian sampling and mixed-model variance estimation rather than model complexity.

DL models implemented in MultiGS-P showed moderate CPU runtimes, with fully connected architectures generally faster than graph-based models, which incurred additional overhead due to graph construction and message passing. Hybrid models exhibited intermediate runtimes and remained computationally feasible for routine use. Across all model classes, PC-based marker representations consistently reduced training time.

GPU acceleration substantially reduced training time for most DL models, with reductions ranging from ∼40% to >90%, but provided little benefit for classical linear and ML models and, in some cases, increased runtime due to data-transfer and workflow overhead. Third-party CNN-based DL models showed markedly longer CPU runtimes than all MultiGS-P models, although GPU acceleration reduced their training time when supported. While absolute runtimes were influenced by system load at the time of benchmarking, relative trends across model classes were consistent, indicating that MultiGS-P achieves practical computational efficiency for large-scale genomic prediction.

## DISCUSSION

### Genomic prediction performance across practical breeding scenarios

The primary objective of MultiGS was not to advocate a single superior genomic prediction model, but rather to provide a unified, practical, decision-support framework for evaluating and deploying multiple GS methodologies under realistic breeding scenarios. Across wheat2000 and maize6000 datasets, where training and test sets were randomly sampled from the same population, DL, hybrid, and ensemble models implemented in MultiGS achieved PAs comparable to RR-BLUP and frequently exceeded those of GBLUP. These results are consistent with previous reports showing that non-linear models can match or marginally improve upon linear mixed models when the training population size is sufficiently large and population structure is well matched between training and testing sets (González-Recio et al. 2014; Montesinos-López et al. 2018).

However, from a breeding perspective, such within-population evaluations reflect an optimistic assessment of prediction performance. Because training and test sets are drawn from the same population, these evaluations closely resemble cross-validation and do not fully capture the challenges of predicting truly new breeding lines in deployment. In contrast, the flax287 dataset provides a realistic across-population prediction case, characterized by a small training population and evaluation in genetically narrow biparental populations. This setting more closely reflects operational breeding programs, where PA often declines sharply due to population divergence, limited training data, and changes in LD patterns (de Los Campos et al. 2013; Habier et al. 2007). For breeders with limited training data or across-population prediction objectives, linear and BLUP-integrated hybrid models are recommended. DL models may be beneficial when training populations are large, marker density is high, and population structure is stable.

The training population size analysis further clarifies these observations. Using the maize6000 dataset, we showed that prediction accuracies increased monotonically with training population size across all model classes, with the largest gains occurring between approximately 500 and 2,500 individuals (**Figure 4**). Notably, linear models such as RR-BLUP continued to improve with increasing sample size and often matched or exceeded deep learning models even at the largest training sizes, highlighting that data availability remains a dominant driver of prediction accuracy regardless of model complexity.

### Model robustness under across-population prediction

The flax results highlight a clear distinction between model capacity and model robustness. Classical linear models, including RR-BLUP, BRR, and Bayesian regressions, provided stable and interpretable baseline performance across traits, particularly for DTM, where prediction accuracy was uniformly low. This stability underscores the continued relevance of linear mixed models in GS, especially for traits dominated by additive genetic effects and for scenarios with limited training data (Meuwissen et al. 2001; VanRaden 2008).

In contrast, many pure DL architectures exhibited high variability and, in some cases, poor or negative predictive performance in flax, reflecting well-known limitations of high-capacity models under small sample sizes and pronounced training–test distribution shifts. These results caution against indiscriminate applications of DL models in breeding programs without careful consideration of population structure and data availability.

Importantly, a subset of DL and hybrid models had improved robustness under this challenging setting. Graph-based models (GraphSAGEGS, GraphFormer) and BLUP-integrated hybrids (DeepBLUP, DeepResBLUP) benefited from stronger inductive biases toward biologically meaningful structure. Graph-based models operate on sample-level genetic relationship graphs rather than raw marker effects, making predictions less sensitive to population-specific LD patterns and allele-frequency shifts. In particular, the inductive neighborhood aggregation in GraphSAGEGS facilitates transfer of information to genetically divergent populations. Similarly, BLUP-integrated hybrids preserve additive genetic effects by anchoring predictions to a linear RR-BLUP component, allowing the deep network to model only residual nonlinear signals. Together, these design choices enhance the stability of DL-based genomic prediction under realistic across-population breeding scenarios.

### Comparison with previously published deep learning models

Although DL models have been applied to GS since 2018 (Montesinos-López et al. 2018), beginning with convolution-based models such as DeepGS (Ma et al. 2018) and have expanded to include recurrent, attention-based, Transformer, and graph-based architectures (Wu et al. 2023; Yan et al. 2025; He et al. 2025; Ma et al. 2024; Wang et al. 2025a; Wang et al. 2023; Ye et al. 2025), most studies report prediction accuracies comparable to, but not consistently exceeding, linear baselines such as RR-BLUP, particularly under large training populations and dense marker settings. Despite methodological advances, many published DL-based GS tools face practical limitations that hinder reproducibility and routine adoption, including incomplete documentation, rigid input requirements, discrepancies between published descriptions and available code, and the lack of standardized workflows for preprocessing and evaluation.

Computational constraints further limit flexibility, as many tools rely on fixed marker dimensions to control runtime and memory usage, requiring ad hoc SNP subsetting or padding (Wang et al. 2025a; Wang et al. 2025b; Deng et al. 2025). In contrast, MultiGS imposes no explicit restrictions on marker number and instead emphasizes biologically informed and computationally efficient representations. Haplotype and PC encodings reduce feature dimensionality while preserving linkage disequilibrium and population structure, improving training efficiency without arbitrary feature selection. Graph-based models in MultiGS further reduce complexity by operating on sample-level graphs rather than marker-level graphs.

Across wheat2000, maize6000, and flax287 datasets, DL and hybrid models implemented in MultiGS achieved prediction accuracies comparable to or exceeding those of previously published DL tools. While third-party models performed well under within-population validation, their performance declined in the flax across-population scenario, whereas several MultiGS hybrid models showed greater stability across datasets and prediction settings. These results suggest that architectural complexity alone is insufficient for robust genomic prediction; instead, models that integrate additive genetic effects with DL refinement and emphasize generalizability are better suited to realistic breeding scenarios. By providing a unified, configurable, and well-documented framework, MultiGS addresses key limitations of existing DL-based GS tools and facilitates fair benchmarking and practical deployment.

### Computational efficiency and breeding deployment

In practical breeding pipelines, computational efficiency is a critical but often underappreciated factor. Most DL and hybrid models implemented in MultiGS required less computational time than previously published DL approaches while delivering comparable predictive accuracy. This efficiency enables frequent model retraining as new phenotypic data becomes available and facilitates large-scale benchmarking across traits, populations, and marker representations.

From an operational perspective, when PA is similar, reduced computational burden becomes a decisive advantage. The ability to execute MultiGS models under CPU-only environments further lowers barriers to adoption, particularly for public breeding programs with limited computational infrastructure. These considerations are essential for translating methodological advances into routine breeding practice.

### Implications for model selection and tool development

Taken together, the results reinforce several key principles for genomic selection. First, no single model is universally optimal across traits or prediction scenarios. Second, classical linear models remain strong and reliable baselines, particularly for across-population prediction. Third, DL models can offer advantages for certain traits and datasets, but their success depends strongly on training population size, genomic architecture, and model design. Hybrid and ensemble approaches consistently provide the most stable improvements, combining the interpretability of linear models with the flexibility of nonlinear learning.

The primary contribution of MultiGS lies in enabling breeders and researchers to explore these trade-offs systematically within a unified framework. By integrating R- and Python-based models, supporting multiple marker representations, and providing standardized evaluation pipelines, MultiGS facilitates informed model selection rather than relying on a single methodology. This design aligns with actual breeding workflows, where adaptability, robustness, and computational practicalities are as important as peak PA.

In addition to model choice, marker representation emerged as a critical and often underappreciated factor influencing prediction accuracy. Across both maize and flax datasets, haplotype-based markers consistently matched or exceeded SNP-based predictions and outperformed PC representations in across-population settings (**Figure 3**). These results indicate that preserving local LD and multi-allelic information can improve robustness and accuracy, particularly when training populations are limited or genetically divergent. Consequently, effective genomic selection requires joint consideration of marker representation, model architecture, and training population characteristics rather than optimization of predictive models alone.

In summary, MultiGS bridges methodological innovation with breeding realities, providing a flexible and efficient platform for deploying genomic selection across diverse crops and prediction scenarios.

### Limitations and future directions

Limitations of this study should be noted. Benchmarking in the wheat2000 and maize6000 datasets relied on random within-population training–test splits, which approximate cross-validation and may overestimate prediction performance relative to actual breeding deployment, particularly for DL models. The flax287 dataset provided a true across-population prediction scenario, but its small training population limited the evaluation of high-capacity DL architectures. Future studies using larger and more diverse populations, combined with systematic across-population and across-environment validations, are warranted to better define conditions under which DL models provide consistent advantages. In addition, the current MultiGS implementation focuses on single-trait prediction, whereas many breeding programs target correlated traits evaluated across multiple environments; extending the framework to multi-trait and multi-environment models should be incorporated in future iterations. Finally, although graph-based and hybrid DL models showed increased robustness in some settings, their performance remained sensitive to marker representation and population structure, highlighting the need for improved hyperparameter optimization, automation, and resource management to support stable and scalable deployment in practical breeding pipelines. Future development will also support matrix-based, scored marker inputs beyond VCF, enabling direct use of breeder-curated genotype tables and alternative marker systems that are not readily convertible to VCF format.

## CONCLUSIONS

MultiGS is designed for both methodological researchers and applied breeding programs, enabling transparent benchmarking as well as routine genomic prediction. MultiGS provides a unified and practical framework for genomic selection by integrating traditional statistical models, ML methods, and modern DL architectures within a standardized workflow. Across wheat2000, maize6000, and flax287 datasets, the results show that while classical linear models such as RR-BLUP remain strong and reliable baselines, selected DL, hybrid, and ensemble models implemented in MultiGS achieve comparable or superior PA under appropriate conditions.

Importantly, the flax across-population case study demonstrates that prediction robustness, rather than peak accuracy under idealized validation, remains the primary challenge for actual breeding applications. In this setting, graph-based and BLUP-integrated hybrid models exhibited more stable generalization than many high-capacity DL architectures. In addition, all MultiGS DL and hybrid models delivered competitive accuracies with lower computational cost than previously published DL tools, supporting their suitability for routine breeding use.

Overall, no single model is universally optimal across traits or populations. MultiGS addresses this fact by providing breeders and researchers with a flexible, efficient, and extensible platform to evaluate and deploy genomic prediction models under genuine breeding scenarios.

## DATA AND CODE AVAILABILITY

All datasets used in this study were obtained from publicly available sources as described in Materials and Methods. The software and various utility programs for this research are freely available on GitHub (https://github.com/AAFC-ORDC-Crop-Bioinfomatics/MultiGS).

## AUTHOR CONTRIBUTIONS

FY: conceptualization, funding acquisition, investigation, methodology, software development, original draft writing. SC: conceptualization, funding acquisition, DNA preparation for SNPs, data curation, review and editing. BT: conceptualization, funding acquisition, data curation, writing–review and editing. JD: software development. CZ, JD. PL, KJ and MH: formal analysis, visualization, data curation, review and editing.

## ACKNOWLEDGMENTS

We thank Dr. Liqian He for reviewing the draft manuscript and providing valuable comments and suggestions.

