## Supplementary Methods, Supplementary figures and tables for "MultiGS: A comprehensive and user-friendly genomic prediction platform Integrating statistical, machine learning, and deep learning models for breeders"

### Supplementary Materials and Methods

#### MultiGS-P

The DL architectures implemented in MultiGS-P are grouped into fully connected, graph-based, hybrid, and BLUP-integrated categories. Each architecture is described in the following sections, with documentation of its design rationale and intended use (Figures S1 and S2). All ML and DL models in MultiGS-P are fully configurable through a centralized configuration file (Table S3), allowing users to adjust hyperparameters, model depth, learning schedules, and regularization settings without modifying source code. This design facilitates systematic benchmarking and fair comparison across diverse model classes while supporting flexible adaptation to different datasets and breeding scenarios.

##### *DNNGS*

Fully connected feedforward neural networks (multilayer perceptrons) have been widely used as baseline DL models for genomic prediction, providing a flexible nonlinear extension of linear mixed models (Montesinos-Lopez et al. 2021). Our DNNGS follows this established MLP paradigm but is implemented as a compact, reproducible architecture with dropout-based regularization and optional batch normalization, and it is designed to accept multiple marker representations (SNPs, haplotypes, or PCs) under a unified MultiGS workflow to enable fair cross-model benchmarking (Figure 1A, Table 1). The model begins with an input-dropout layer followed by four sequential fully connected blocks with hidden dimensions 512, 256, 128, and 64. Each block consists of a dense layer with ReLU activation, dropout, and optional batch normalization to improve stability. A final dense output layer generates trait predictions. The architecture provides a balance between modeling nonlinear genotype–phenotype relationships and maintaining computational efficiency.

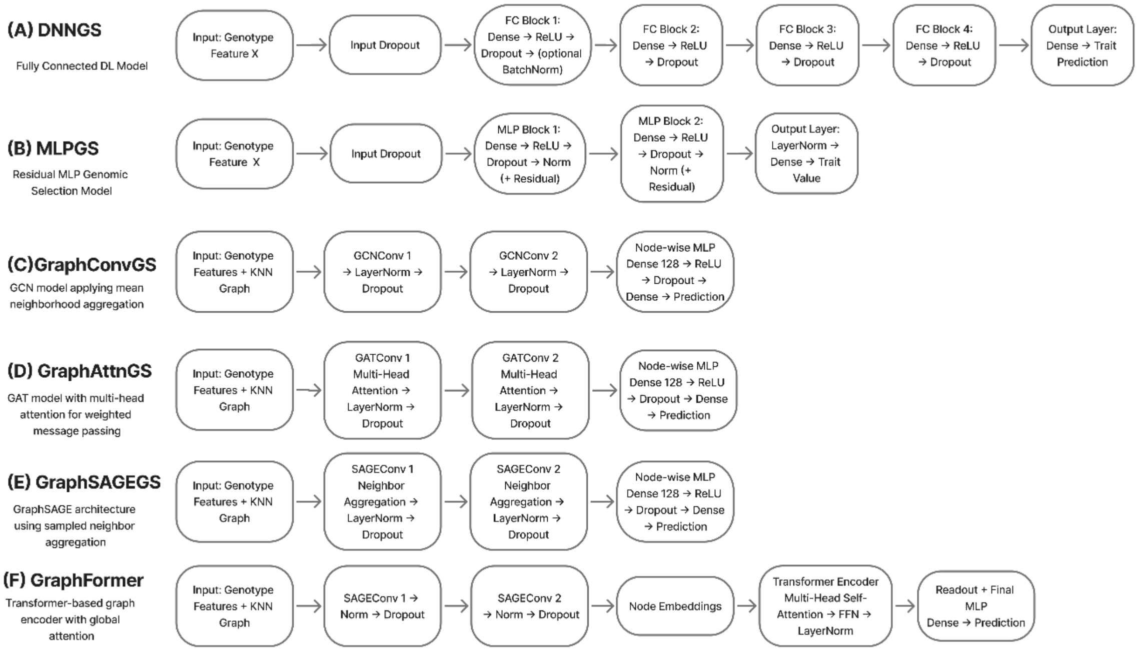


**Figure S1.** Architectures of two fully connected network and four graph-based deep learning models for genomic selection: (**A**) DNNGS, (**B**) MLPGS, (**C**) GraphConvGS, (**D**) GraphAttnGS, (**E**) GraphSAGEGS, and (**F**) GraphFormer.

***MLPGS***

MLPGS is a multilayer perceptron architecture incorporating normalization and optional residual connections to enable stable training on genomic features (Figure 1B, Table 1). The model applies input dropout followed by two fully connected blocks: the first typically includes a 128-unit dense layer with GELU or ReLU activation, dropout, and LayerNorm or BatchNorm; the second block (64 units) follows the same structure. Optional residual connections allow block inputs to be added to outputs when dimensions match. A final normalization step and dense output layer produce trait predictions. This design offers improved regularization and gradient stability compared to conventional MLPs. While MLPs have been previously used as nonlinear genomic prediction models, our MLPGS variant emphasizes training stability and regularization through explicit normalization and residual connections, and its implementation within MultiGS enables fair, reproducible comparison with other model families.

***GraphConvGS***

Graph neural networks (GNNs) have recently been introduced for genomic prediction to explicitly model genetic relationships among individuals by representing samples as nodes connected through similarity-based edges (Kihlman et al. 2024; He et al. 2025). GraphConvGS follows this paradigm by constructing sample-level k-nearest-neighbor genotype graphs (convolutional neural network, GCN) and applying graph convolution to aggregate information from genetically similar individuals (Figure 1C, Table 1). Individuals are represented as nodes with marker-derived features, and edges encode genetic similarity. The architecture includes two GCNConv layers, each followed by LayerNorm, ReLU activation, and dropout, enabling neighborhood aggregation across genetically similar samples. A node-wise MLP (Dense → ReLU → Dropout → Dense) is applied to the resulting node embeddings to produce trait predictions. GraphConvGS captures relational patterns between individuals that are not available to feature-only models.

***GraphAttnGS***

Graph attention networks (GATs) were introduced by Veličković et al. (2018) as an extension of GCNs that learn attention weights over neighbors rather than using uniform or degree-normalized aggregation. GATs are now widely used in biological and population-structure problems where neighbor importance is heterogeneous, however few GS studies explored attention-based graph models (He et al. 2025) to address variable genetic similarity and population structure. GraphAttnGS extends GraphConvGS by replacing its convolution layers with GATConv layers, enabling multi-head attention over graph neighbors learn which neighbors matter most, stabilizing training and capturing multiple "views" of genetic similarity simultaneously (Figure 1D, Table 1). Two stacked GATConv layers (each with normalization, activation, and dropout) learn node embeddings by weighting neighbor contributions according to learned attention coefficients. A node-wise MLP produces final predictions. This architecture adaptively highlights the most informative neighbors and models heterogeneous genotype similarity patterns across individuals.

***GraphSAGEGS***

Unlike transductive GCN- and GAT-based models, GraphSAGEGS summarizes local neighborhood information through learned aggregation functions, providing robust performance when predicting genetically distinct or previously unseen populations (Figure 1E, Table 1). The architecture includes two SAGEConv layers, each followed by normalization, activation, and dropout. A node-level MLP then maps these embeddings to predicted trait values. By aggregating summary statistics from each node's local neighborhood, GraphSAGEGS provides efficient and robust performance, especially in across-population prediction scenarios.

***GraphFormer***

Hybrid graph–Transformer architectures have recently emerged as an effective strategy for combining local message passing with global self-attention, enabling simultaneous modeling of neighborhood structure and long-range dependencies (Gao et al. 2023; Ramezankhani et al. 2025). However, few genomic selection pipelines integrate both components within a unified and systematically benchmarked framework. GraphFormer adopts this strategy by combining GraphSAGE-based local aggregation with Transformer-style global attention across individuals. Specifically, two SAGEConv layers are first applied to generate node embeddings that capture local genetic neighborhoods. These embeddings are then processed by a Transformer encoder, typically comprising two layers with multi-head self-attention and feed-forward networks, to model global interactions among all individuals. A readout module followed by a final MLP produces trait predictions. By explicitly separating local relational learning from global interaction modeling, GraphFormer captures both fine-scale genetic structure and long-range population-level dependencies within the population graph (Figure 1F, Table 1).

***DeepResBLUP***

Hybrid strategies that augment BLUP predictions with nonlinear deep-learning components have been explored such as DLGBLUP (Shokor et al. 2025), to account for genetic effects beyond linear additive assumptions. Building on this concept, DeepResBLUP is designed as a residual learning framework that explicitly models nonlinear deviations from a classical RR-BLUP baseline (Figure 2A, Table 1). In DeepResBLUP, RR-BLUP is first fitted to the genotype matrix to generate baseline GEBVs, which are treated as a strong additive prior. These RR-BLUP predictions are then provided as fixed, or optionally weakly trainable, inputs to a deep neural network that is constrained to learn residual corrections rather than replace the linear model. The residual network consists of three fully connected layers (256 → 128 → 64 units) with GELU activation, batch normalization, and dropout. Its output represents the nonlinear residual component, which is combined with the original RR-BLUP prediction through an explicit skip connection to produce the final predictions.

By focusing the deep network on residual signals, DeepResBLUP preserves the interpretability and robustness of RR-BLUP while selectively capturing nonlinear effects not explained by additive marker effects. In addition, the framework provides flexibility by allowing RR-BLUP to be replaced with alternative linear models, enabling residual learning on top of different additive baselines within the same architecture.

***DeepBLUP***

Recent work has shown that BLUP-style components can be made differentiable and jointly trained with neural networks (Guhlin and Dearden 2025); DeepBLUP operationalizes this idea for RR-BLUP by embedding an RR-BLUP-initialized linear layer within an end-to-end architecture, with options to fix or fine-tune the BLUP layer and to enable a stabilizing skip connection. DeepBLUP integrates RR-BLUP directly into a fully end-to-end trainable neural architecture by implementing it as the first linear layer of the network (Figure 2B, Table 1). This RR-BLUP-initialized layer maps marker effects to predictions and can be either fixed or jointly optimized during training. Unlike DeepResBLUP, RR-BLUP in DeepBLUP is not treated as a standalone baseline but rather as an embedded component within the network. The RR-BLUP layer feeds into a sequence of dense layers (256 → 128 → 64 units) with GELU activation, batch normalization, and dropout, allowing nonlinear feature transformations to be learned directly from the RR-BLUP output. A final dense layer produces the predictions. An optional skip connection may be enabled to stabilize training by adding the RR-BLUP output to the network's final predictions, but the model is fundamentally optimized as a single unified system, rather than as a baseline-plus-residual model.

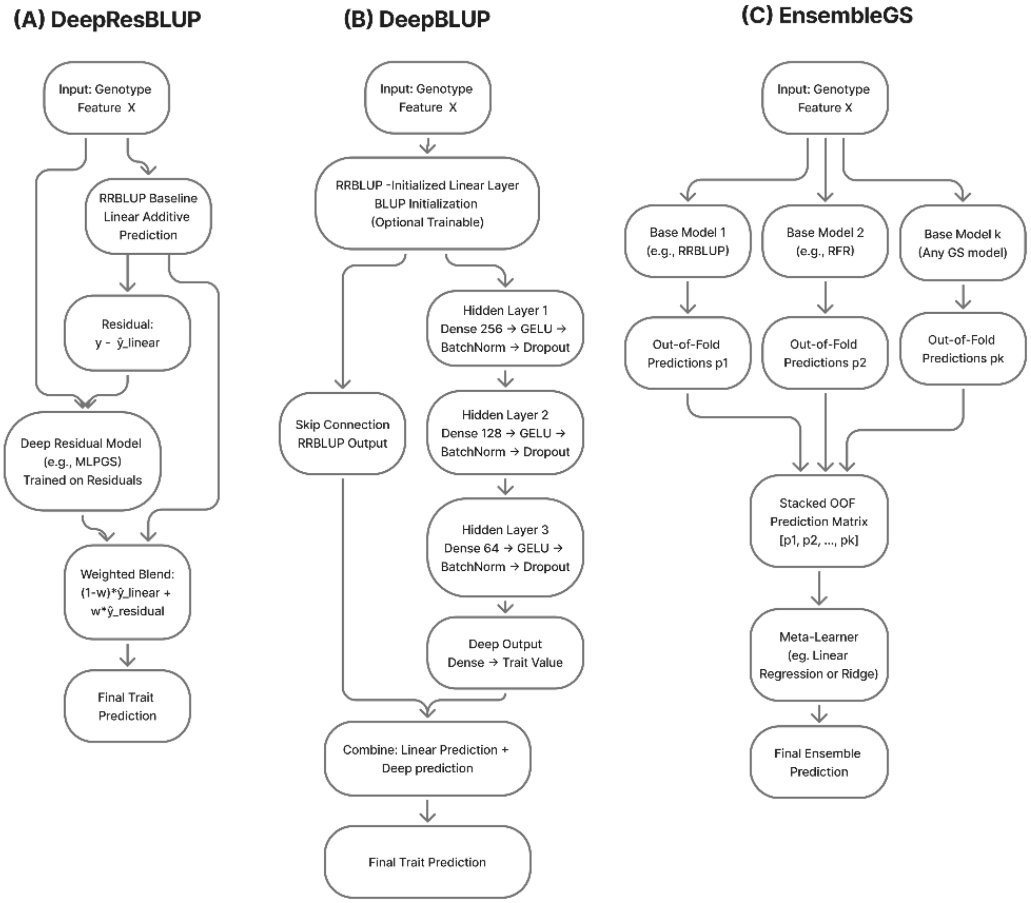


**Figure S2. Architectures of three hybrid genomic selection models that integrate linear and deep learning components**. **(A)** DeepResBLUP, **(B)** DeepBLUP and **(C)** EnsembleGS.

***EnsembleGS***

Stacking-based ensemble learning has previously been applied to genomic prediction to improve accuracy and robustness by optimally combining diverse learners using out-of-fold predictions (Nascimento et al. 2024; Liang et al. 2021). EnsembleGS extends these approaches by supporting stacking over arbitrary MultiGS models, including linear, ML, DL, and hybrid architectures, within a standardized preprocessing and evaluation workflow (Figure 2C, Table 1). Unlike many prior implementations that stack a fixed set of learners, EnsembleGS allows users to flexibly configure both the base-model library and the meta-learner through the MultiGS configuration system.

In EnsembleGS, a set of independent base models (e.g., RR-BLUP, BRR, XGBoost, LightGBM, and DNNGS) implemented in MultiGS-P are trained to generate out-of-fold (OOF) predictions, which are concatenated into a stacked prediction matrix. A meta-learner—by default linear regression, though alternative learners are supported—is then trained on this matrix to produce final predictions. During inference, predictions from the trained base models are passed through the meta-learner to yield the ensemble output. By leveraging complementary strengths across linear, ML, and DL models, EnsembleGS typically provides improved stability and robustness of prediction performance across traits and datasets, consistent with previous stacking ensemble applications in genomic prediction.

### Supplementary Figures and Tables

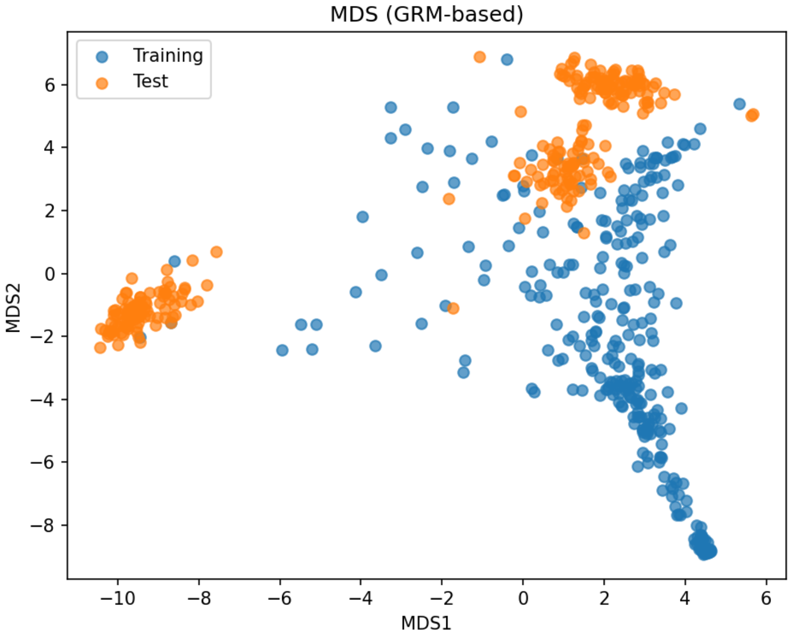


**Figure S3.** Multidimensional scaling (MDS) analysis based on the genomic relationship matrix (GRM) for 278 training lines (flax287) and 260 test lines from three biparental populations, showing pronounced genetic structure between the two sets.

**Table S1.** Linear and machine learning models implemented in MultiGS-R.

| Model name | Full name | Model category | Core algorithm | Key features | R package(s) |
|---|---|---|---|---|---|
| **RR-BLUP** | Ridge Regression Best Linear Unbiased Prediction | Linear mixed model | Penalized linear regression with ridge penalty | Assumes equal variance of marker effects; computationally efficient baseline | *rrBLUP* |
| **GBLUP** | Genomic Best Linear Unbiased Prediction | Linear mixed model | Genomic relationship matrix–based BLUP | Models additive genetic relationships using genomic kinship | *BGLR* |
| **BRR** | Bayesian Ridge Regression | Bayesian linear model | Bayesian ridge regression | Shrinkage of marker effects with Gaussian prior | *BGLR* |
| **BL** | Bayesian LASSO | Bayesian linear model | LASSO with Laplace prior | Allows variable shrinkage across markers | *BGLR* |
| **BayesA** | BayesA | Bayesian linear model | Marker-specific variance model | Heavy-tailed priors capture large-effect loci | *BGLR* |
| **BayesB** | BayesB | Bayesian linear model | Mixture model with spike-and-slab prior | Performs variable selection by excluding many markers | *BGLR* |
| **BayesC** | BayesC | Bayesian linear model | Modified BayesB with shared variance | Improved stability and reduced sensitivity to hyperparameters | *BGLR* |
| **RKHS** | Reproducing Kernel Hilbert Space regression | Kernel-based model/ML | Gaussian kernel regression | Captures nonlinear and epistatic effects | *BGLR* |

| Model name | Full name | Model category | Core algorithm | Key features | R package(s) |
|---|---|---|---|---|---|
| **RFR** | Random Forest Regression | ML | Ensemble of decision trees | Captures nonlinear interactions; robust to noise | *randomForest* |
| **RFC** | Random Forest Classification | ML | Ensemble of decision trees | Used for categorical trait prediction | *randomForest* |
| **SVR** | Support Vector Regression | ML | Kernel-based margin regression | Effective in high-dimensional settings | *e1071* |
| **SVC** | Support Vector Classification | ML | Kernel-based classification | Used for binary or multiclass traits | *e1071* |

*rrBLUP*: Endelman (2011); *BGLR*: Perez and de los Campos (2014); *randomForest*: Liaw and Wiener (2002); *e1017*: Meyer et al. (2023).

**Table S2.** Summary of eight linear and machine learning models implemented in MultiGS-P.

| Model | Architecture / Type | Core Algorithm / Method | Key Features | Best Use Cases |
|---|---|---|---|---|
| **R_RRBLUP** | Linear Mixed Model (R) | Ridge regression BLUP via R package *rrBLUP* | Widely validated baseline | Additive traits |
| **R_GBLUP** | Linear Mixed Model (R) | Genomic relationship kernel BLUP | Captures population structure | Standard GS benchmark |
| **RRBLUP** | Linear regression (Python) | Ridge regression | Consistent with R version | Additive effects |
| **ElasticNet** | Linear model (L1+L2) | Elastic-net regularization | Feature shrinkage | Sparse/noisy SNP effects |
| **BRR** | Bayesian linear regression | Gaussian priors | Uncertainty estimation | Moderate shrinkage traits |
| **RFR** | Ensemble of trees | Random Forest | Nonlinear interactions | Epistasis / nonlinear |
| **XGBoost** | Gradient boosting trees | Additive boosting | Handles complex patterns | Large SNP sets |
| **LightGBM** | Gradient boosting trees | Histogram-based boosting | Fast, scalable | High-dimensional SNPs |

**Table S3**: Default hyperparameter settings for the machine learning and deep learning models implemented in MultiGS-P

```
[Hyperparameters_R_RRBLUP]
method = REML    #REML|ML

[Hyperparameters_R_GBLUP]
method = REML    #REML|ML

[Hyperparameters_RRBLUP]
lambda_value = None
method = mixed_model
lambda_method = auto    # Options: auto|reml|heritability|fixed
tol = 1e-8

[Hyperparameters_ElasticNet]
# Reduce regularization for ElasticNet: fro, 1 to 0.1->0.01->0.001
alpha = 1.0
l1_ratio = 0.1   # toward ridge: from 0.5 to 0.1-0.3

[Hyperparameters_LASSO]
alpha = 1.0

[Hyperparameters_XGBoost]
n_estimators = 100
max_depth = 6
learning_rate = 0.1
subsample = 0.8
colsample_bytree = 0.8
random_state = 42

[Hyperparameters_LightGBM]
n_estimators = 100
max_depth = -1
learning_rate = 0.1
num_leaves = 31
subsample = 0.8
colsample_bytree = 0.8
random_state = 42

[Hyperparameters_MLPGS]
hidden_layers = 1024,512,256
activation = gelu
norm = layer
residual = true
input_dropout = 0.05
dropout = 0.5
```

```
learning_rate = 0.0005
weight_decay = 0.0015
batch_size = 16
epochs = 300
early_stopping_patience = 20
warmup_ratio = 0.1
grad_clip = 1.0
seeds = 3
use_huber = true
huber_delta = 1.0
swa = true
swa_start = 0.7
swa_freq = 1

[Hyperparameters_DNNGS]
hidden_layers = 512,256,128,64
learning_rate = 0.001
batch_size = 32
epochs = 300
dropout = 0.3
activation = gelu
batch_norm = true
weight_decay = 0.0001
input_dropout = 0.1

[Hyperparameters_GraphConvGS]
hidden_channels = 128
num_layers = 2
hidden_mlp = 128
dropout = 0.2
learning_rate = 0.0005
epochs = 500
top_k = 20
graph_method = knn
knn_metric = euclidean
patience = 20

[Hyperparameters_GraphAttnGS]
hidden_channels = 128
num_layers = 2
heads = 4
hidden_mlp = 128
dropout = 0.2
learning_rate = 0.0005
epochs = 500
top_k = 20
graph_method = knn
knn_metric = euclidean
```

patience = 20

[Hyperparameters_GraphSAGEGS]
hidden_channels = 128
num_layers = 2
hidden_mlp = 128
dropout = 0.2
learning_rate = 0.0005
epochs = 500
top_k = 20
graph_method = knn
knn_metric = euclidean
aggr = mean
patience = 20

[Hyperparameters_GraphFormer]
#GraphFormer:
gnn_type = SAGE # Choose: SAGE|GraphConvGS|GraphAttnGS
gnn_hidden = 128 # Output size of GNN layer
transformer_layers = 2 # Number of transformer layers
d_model = 128 # Transformer dimension
nhead = 4 # Number of attention heads
mlp_hidden = 128 # MLPGS hidden size
learning_rate = 0.001
epochs = 500
patience = 30
dropout = 0.1
weight_decay = 0.001
graph_method = knn
knn_metric = euclidean
top_k = 30

[Hyperparameters_DeepResBLUP]
base_model = R_RRBLUP
dl_model = MLPGS # MLPGS|HybridAttnMLPGS|DNNGS|AttnCNNGS|hybrid (hybrid: marker-transformer + optional sample GNN, very slow)
dl_hidden_layers = 128,64
dl_dropout = 0.2
dl_learning_rate = 0.001
dl_batch_size = 32
dl_epochs = 100

[Hyperparameters_DeepBLUP]
rrblup_lambda = 0.001
dl_hidden_layers = 128,64,32
dropout = 0.3
activation = gelu
use_precomputed_rrblup = true

```
train_rrblup_layer = true
learning_rate = 0.0001
batch_size = 16
epochs = 200
weight_decay = 0.0001
use_batch_norm = true
use_residual_connections = true

[Hyperparameters_EnsembleGS]
# models available for stacking
# 'R_RRBLUP', 'R_GBLUP', 'RRBLUP',
# 'ElasticNet', 'RFR', 'BRR',
# 'XGBoost', 'LightGBM',
# 'DNNGS', 'AttnCNNGS', 'MLPGS','HybridAttnMLPGS',
# 'GraphConvGS','GraphAttnGS', 'GraphSAGEGS',
# 'GraphFormer', 'Transformer',
# 'DeepResBLUP', 'DeepBLUP',
base_models = R_RRBLUP,ElasticNet,LightGBM, MLPGS, GraphSAGEGS
meta_model = linear   # linear|ridge
meta_alpha = 1.0
```

**Table S4.** Genetic diversity and population differentiation between training and test sets across for three datasets

| Dataset | Population | Nucleotide diversity (π) | Heterozygosity (Ho) | Number of SNPs | Number of individuals | FST (training vs. test ) |
|---|---|---|---|---|---|---|
| Wheat2000 | Training | 0.1353 | 0.0189 | 9,927 | 4,000 | $-9.14 \times 10^{-6}$ |
| | Test | 0.1340 | 0.0184 | 9,927 | 1,600 | |
| Maize6000 | Training | 0.3003 | 0.3811 | 10,000 | 4,664 | -0.001 |
| | Test | 0.2990 | 0.3795 | 10,000 | 1,167 | |
| Flax287 | Training | 0.3850 | 0.0142 | 33,596 | 298 | |
| | Test | 0.3716 | 0.0062 | 33,596 | 260 | 0.2666 |

FST: Fixation index.

**Table S5**. Prediction accuracies of five traits across models implemented in MultiGS-P, evaluated using a wheat training set of 1,600 accessions and a testing set of 400 randomly selected accessions genotyped with a randomly selected set of 10,000 SNP markers.

| **Model** | **Model type** | **Tool** | **Grain hardnes (GH)** | | **Grain length (GL)** | | **Grain protein (GP)** | | **Grain width (GW)** | | **Thousand-kernel weight (TKW)** | |
|---|---|---|---|---|---|---|---|---|---|---|---|---|
| | | | **SNP** | **PC** | **SNP** | **PC** | **SNP** | **PC** | **SNP** | **PC** | **SNP** | **PC** |
| RR-BLUP (R) | Linear mixed | MultiGS-R | 0.584 | 0.581 | 0.725 | 0.721 | 0.500 | 0.510 | 0.743 | 0.739 | 0.657 | 0.644 |
| GBLUP (R) | Linear mixed | MultiGS-R | 0.587 | 0.485 | 0.720 | 0.679 | 0.504 | 0.469 | 0.742 | 0.679 | 0.647 | 0.612 |
| BRR (R) | Bayesian linear | MultiGS-R | 0.586 | 0.487 | 0.716 | 0.678 | 0.507 | 0.469 | 0.743 | 0.677 | 0.643 | 0.614 |
| BL (R) | Bayesian linear | MultiGS-R | 0.588 | 0.550 | 0.716 | 0.709 | 0.504 | 0.481 | 0.746 | 0.721 | 0.644 | 0.641 |
| BayesA (R) | Bayesian linear | MultiGS-R | 0.585 | 0.555 | 0.719 | 0.709 | 0.505 | 0.478 | 0.747 | 0.722 | 0.645 | 0.641 |
| BayesB (R) | Bayesian linear | MultiGS-R | 0.588 | 0.543 | 0.716 | 0.693 | 0.500 | 0.475 | 0.747 | 0.712 | 0.644 | 0.621 |
| BayesC (R) | Bayesian linear | MultiGS-R | 0.587 | 0.518 | 0.716 | 0.645 | 0.504 | 0.473 | 0.741 | 0.677 | 0.644 | 0.594 |
| RFR (R) | Machine learning | MultiGS-R | 0.613 | 0.569 | 0.743 | 0.731 | 0.528 | 0.533 | 0.757 | 0.740 | 0.668 | 0.665 |
| SVR (R) | Machine learning | MultiGS-R | 0.513 | 0.470 | 0.650 | 0.657 | 0.416 | 0.454 | 0.679 | 0.683 | 0.569 | 0.566 |
| RKHS (R) | Kernel-based/Machine learning | MultiGS-R | 0.585 | 0.500 | 0.715 | 0.691 | 0.506 | 0.465 | 0.738 | 0.655 | 0.656 | 0.604 |
| RFC (R) | Machine learning | MultiGS-R | 0.595 | 0.542 | 0.671 | 0.662 | 0.491 | 0.473 | 0.611 | 0.563 | 0.589 | 0.573 |

| Model | Model type | Tool | Grain hardnes (GH) | | Grain length (GL) | | Grain protein (GP) | | Grain width (GW) | | Thousand-kernel weight (TKW) | |
|---|---|---|---|---|---|---|---|---|---|---|---|---|
| | | | SNP | PC | SNP | PC | SNP | PC | SNP | PC | SNP | PC |
| SVC (R) | Machine learning | MultiGS-R | 0.498 | 0.410 | 0.638 | 0.601 | 0.430 | 0.442 | 0.558 | 0.495 | 0.553 | 0.562 |
| R_RRBLUP | Linear mixed | MultiGS-P | 0.586 | 0.592 | 0.717 | 0.710 | 0.504 | 0.505 | 0.739 | 0.736 | 0.644 | 0.638 |
| R_GBLUP | Linear mixed | MultiGS-P | 0.183 | 0.183 | 0.153 | 0.151 | 0.099 | 0.097 | 0.237 | 0.237 | 0.140 | 0.139 |
| RRBLUP | Linear mixed | MultiGS-P | 0.582 | 0.588 | 0.716 | 0.700 | 0.491 | 0.482 | 0.740 | 0.723 | 0.640 | 0.615 |
| ElasticNet | Linear | MultiGS-P | 0.477 | 0.518 | 0.617 | 0.657 | 0.386 | 0.459 | 0.683 | 0.698 | 0.561 | 0.606 |
| BRR | Bayesian linear regression | MultiGS-P | 0.586 | 0.592 | 0.717 | 0.710 | 0.504 | 0.505 | 0.739 | 0.736 | 0.644 | 0.638 |
| RFR | Ensemble of trees | MultiGS-P | 0.573 | 0.540 | 0.726 | 0.730 | 0.529 | 0.529 | 0.732 | 0.736 | 0.656 | 0.660 |
| XGBoost | Gradient boosting trees | MultiGS-P | 0.605 | 0.577 | 0.721 | 0.730 | 0.462 | 0.514 | 0.743 | 0.721 | 0.639 | 0.646 |
| LightGBM | Gradient boosting trees | MultiGS-P | 0.623 | 0.565 | 0.726 | 0.721 | 0.471 | 0.486 | 0.740 | 0.716 | 0.646 | 0.633 |
| DNNGS | Deep learning | MultiGS-P | 0.537 | 0.541 | 0.722 | 0.716 | 0.496 | 0.473 | 0.722 | 0.715 | 0.655 | 0.615 |
| HybridAttnMLP | Deep learning | MultiGS-P | 0.523 | 0.542 | 0.703 | 0.685 | 0.443 | 0.493 | 0.713 | 0.702 | 0.640 | 0.635 |
| MLPGS | Deep learning | MultiGS-P | 0.494 | 0.559 | 0.671 | 0.716 | 0.406 | 0.512 | 0.681 | 0.747 | 0.608 | 0.650 |
| GraphConvGS | Deep learning | MultiGS-P | 0.501 | 0.468 | 0.553 | 0.497 | 0.404 | 0.382 | 0.624 | 0.591 | 0.542 | 0.409 |

| **Model** | **Model type** | **Tool** | **Grain hardnes (GH)** | | **Grain length (GL)** | | **Grain protein (GP)** | | **Grain width (GW)** | | **Thousand-kernel weight (TKW)** | |
|---|---|---|---|---|---|---|---|---|---|---|---|---|
| | | | **SNP** | **PC** | **SNP** | **PC** | **SNP** | **PC** | **SNP** | **PC** | **SNP** | **PC** |
| GraphAttnGS | Deep learning | MultiGS-P | 0.425 | 0.465 | 0.564 | 0.596 | 0.341 | 0.395 | 0.601 | 0.602 | 0.497 | 0.495 |
| GraphSAGEGS | Deep learning | MultiGS-P | 0.544 | 0.589 | 0.696 | 0.659 | 0.450 | 0.453 | 0.706 | 0.712 | 0.632 | 0.608 |
| GraphFormer | Deep learning | MultiGS-P | 0.502 | 0.570 | 0.685 | 0.685 | 0.446 | 0.497 | 0.712 | 0.697 | 0.603 | 0.604 |
| DeepResBLUP | Deep learning | MultiGS-P | 0.582 | 0.592 | 0.718 | 0.712 | 0.498 | 0.499 | 0.738 | 0.744 | 0.644 | 0.634 |
| DeepBLUP | Deep learning | MultiGS-P | 0.538 | 0.594 | 0.701 | 0.682 | 0.448 | 0.481 | 0.722 | 0.716 | 0.631 | 0.620 |
| EnsembleGS | Deep learning | MultiGS-P | 0.623 | 0.569 | 0.731 | 0.718 | 0.482 | 0.490 | 0.737 | 0.723 | 0.655 | 0.631 |
| DeepGS | Deep learning | Third-pary | 0.629 | NA | 0.727 | NA | 0.523 | NA | 0.722 | NA | 0.669 | NA |
| CropFormer | Deep learning | Third-pary | 0.510 | NA | 0.706 | NA | 0.389 | NA | 0.703 | NA | 0.640 | NA |
| WheatGP | Deep learning | Third-pary | 0.523 | NA | 0.697 | NA | 0.013 | NA | 0.696 | NA | 0.642 | NA |

**Table S6.** Prediction accuracies of three traits across models implemented in MultiGS-P, evaluated using a maize training set of 4,664 lines and a testing set of 1,167 randomly selected lines, and genotyped with 10,000 randomly selected single nucleotide polymorphism (SNP), 5,439 haplotype (HAP) or 313 principal component (PC) markers.

| **Model** | **Model Type** | **Tool** | **Days to tassel (DTT)** | | | **Ear weight (EW)** | | | **Plant height (PH)** | | |
|---|---|---|---|---|---|---|---|---|---|---|---|
| | | | **SNP** | **HAP** | **PC** | **SNP** | **HAP** | **PC** | **SNP** | **HAP** | **PC** |
| RR-BLUP (R) | Linear mixed | MultiGS-R | 0.934 | 0.930 | 0.918 | 0.764 | 0.759 | 0.721 | 0.925 | 0.923 | 0.878 |
| GBLUP (R) | Linear mixed | MultiGS-R | 0.935 | 0.931 | 0.917 | 0.769 | 0.762 | 0.721 | 0.926 | 0.924 | 0.879 |
| BRR (R) | Bayesian linear | MultiGS-R | 0.935 | 0.931 | 0.917 | 0.768 | 0.762 | 0.721 | 0.927 | 0.924 | 0.879 |
| BL (R) | Bayesian linear | MultiGS-R | 0.936 | 0.932 | 0.918 | 0.771 | 0.765 | 0.722 | 0.928 | 0.926 | 0.879 |
| BayesA (R) | Bayesian linear | MultiGS-R | 0.936 | 0.933 | 0.918 | 0.772 | 0.767 | 0.722 | 0.929 | 0.926 | 0.878 |
| BayesB (R) | Bayesian linear | MultiGS-R | 0.936 | 0.933 | 0.917 | 0.774 | 0.768 | 0.722 | 0.928 | 0.927 | 0.879 |
| BayesC (R) | Bayesian linear | MultiGS-R | 0.935 | 0.932 | 0.916 | 0.773 | 0.764 | 0.721 | 0.927 | 0.927 | 0.879 |
| RFR (R) | Machine learning | MultiGS-R | 0.924 | 0.921 | 0.858 | 0.756 | 0.746 | 0.652 | 0.901 | 0.900 | 0.822 |
| SVR (R) | Machine learning | MultiGS-R | 0.927 | 0.925 | 0.916 | 0.739 | 0.742 | 0.704 | 0.919 | 0.920 | 0.876 |
| RKHS (R) | Kernel-based/Machine learning | MultiGS-R | 0.936 | 0.934 | 0.932 | 0.777 | 0.776 | 0.775 | 0.928 | 0.927 | 0.923 |
| RFC (R) | Machine learning | MultiGS-R | 0.799 | 0.802 | 0.725 | 0.658 | 0.650 | 0.625 | 0.798 | 0.798 | 0.748 |
| SVC (R) | Machine learning | MultiGS-R | 0.796 | 0.794 | 0.782 | 0.655 | 0.664 | 0.655 | 0.795 | 0.795 | 0.761 |

| Model | Model Type | Tool | Days to tassel (DTT) | | | Ear weight (EW) | | | Plant height (PH) | | |
|---|---|---|---|---|---|---|---|---|---|---|---|
| | | | SNP | HAP | PC | SNP | HAP | PC | SNP | HAP | PC |
| R_RRBLUP | Linear mixed | MultiGS-P | 0.935 | 0.931 | 0.915 | 0.768 | 0.762 | 0.715 | 0.926 | 0.924 | 0.873 |
| R_GBLUP | Linear mixed | MultiGS-P | 0.612 | 0.616 | 0.612 | 0.404 | 0.430 | 0.403 | 0.567 | 0.586 | 0.566 |
| RRBLUP | Linear mixed | MultiGS-P | 0.935 | 0.932 | 0.915 | 0.768 | 0.762 | 0.716 | 0.923 | 0.922 | 0.874 |
| ElasticNet | Linear | MultiGS-P | 0.826 | 0.824 | 0.861 | 0.615 | 0.587 | 0.630 | 0.766 | 0.758 | 0.800 |
| BRR | Bayesian linear regression | MultiGS-P | 0.935 | 0.931 | 0.915 | 0.768 | 0.762 | 0.715 | 0.926 | 0.924 | 0.873 |
| RFR | Ensemble of trees | MultiGS-P | 0.904 | 0.904 | 0.834 | 0.726 | 0.720 | 0.617 | 0.883 | 0.879 | 0.808 |
| XGBoost | Gradient boosting trees | MultiGS-P | 0.937 | 0.935 | 0.871 | 0.788 | 0.785 | 0.666 | 0.925 | 0.929 | 0.843 |
| LightGBM | Gradient boosting trees | MultiGS-P | 0.937 | 0.933 | 0.879 | 0.791 | 0.784 | 0.675 | 0.929 | 0.929 | 0.845 |
| DNNGS | Deep learning | MultiGS-P | 0.927 | 0.932 | 0.922 | 0.753 | 0.763 | 0.726 | 0.912 | 0.919 | 0.897 |
| HybridAttnMLP | Deep learning | MultiGS-P | 0.925 | 0.932 | 0.925 | 0.740 | 0.760 | 0.748 | 0.900 | 0.919 | 0.908 |
| MLPGS | Deep learning | MultiGS-P | 0.902 | 0.913 | 0.914 | 0.709 | 0.744 | 0.730 | 0.871 | 0.898 | 0.890 |
| GraphConvGS | Deep learning | MultiGS-P | 0.843 | 0.840 | 0.857 | 0.599 | 0.591 | 0.600 | 0.773 | 0.760 | 0.785 |
| GraphAttnGS | Deep learning | MultiGS-P | 0.819 | 0.844 | 0.811 | 0.517 | 0.568 | 0.523 | 0.739 | 0.730 | 0.740 |
| GraphSAGEGS | Deep learning | MultiGS-P | 0.917 | 0.908 | 0.913 | 0.740 | 0.752 | 0.726 | 0.909 | 0.909 | 0.881 |
| GraphFormer | Deep learning | MultiGS-P | 0.920 | 0.918 | 0.911 | 0.741 | 0.746 | 0.723 | 0.909 | 0.910 | 0.886 |
| DeepResBLUP | Deep learning | MultiGS-P | 0.934 | 0.932 | 0.925 | 0.766 | 0.762 | 0.747 | 0.924 | 0.923 | 0.900 |

| Model | Model Type | Tool | Days to tassel (DTT) | | | Ear weight (EW) | | | Plant height (PH) | | |
|---|---|---|---|---|---|---|---|---|---|---|---|
| | | | SNP | HAP | PC | SNP | HAP | PC | SNP | HAP | PC |
| DeepBLUP | Deep learning | MultiGS-P | 0.935 | 0.929 | 0.926 | 0.768 | 0.758 | 0.765 | 0.924 | 0.917 | 0.908 |
| EnsembleGS | Deep learning | MultiGS-P | 0.919 | 0.903 | 0.901 | 0.745 | 0.752 | 0.720 | 0.914 | 0.911 | 0.881 |
| DeepGS | Deep learning | Third-party | 0.934 | NA | NA | 0.764 | NA | NA | 0.925 | NA | NA |
| CropFormer | Deep learning | Third-party | 0.914 | NA | NA | 0.692 | NA | NA | 0.898 | NA | NA |
| DPCFormer | Deep learning | Third-party | 0.892 | NA | NA | 0.686 | NA | NA | 0.843 | NA | NA |
| WheatGP | Deep learning | Third-party | 0.843 | NA | NA | 0.765 | NA | NA | 0.923 | NA | NA |

**Table S7.** Prediction accuracies of three traits across models implemented in MultiGS, evaluated using a flax training set of 278 accessions from a core collection and a testing set of 260 biparental inbred lines, with 7,363 haplotype markers derived from 33,895 common SNPs.

| **Model** | **Model type** | **Tool** | **Days to maturity (DTM)** | | | **Oil content (OIL)** | | | **Plant height (PH)** | | |
|---|---|---|---|---|---|---|---|---|---|---|---|
| | | | **SNP** | **HAP** | **PC** | **SNP** | **HAP** | **PC** | **SNP** | **HAP** | **PC** |
| RR-BLUP (R) | Linear mixed | MultiGS-R | 0.359 | 0.367 | 0.372 | 0.508 | 0.661 | 0.498 | 0.540 | 0.590 | 0.553 |
| GBLUP (R) | Linear mixed | MultiGS-R | 0.325 | 0.343 | 0.047 | 0.450 | 0.596 | 0.095 | 0.553 | 0.605 | -0.058 |
| BRR (R) | Bayesian linear | MultiGS-R | 0.336 | 0.383 | 0.063 | 0.495 | 0.661 | 0.089 | 0.540 | 0.595 | -0.072 |
| BL (R) | Bayesian linear | MultiGS-R | 0.350 | 0.361 | 0.043 | 0.436 | 0.575 | 0.047 | 0.613 | 0.635 | 0.306 |
| BayesA (R) | Bayesian linear | MultiGS-R | 0.336 | 0.361 | 0.035 | 0.483 | 0.615 | 0.006 | 0.541 | 0.626 | 0.393 |
| BayesB (R) | Bayesian linear | MultiGS-R | 0.360 | 0.372 | 0.028 | 0.487 | 0.660 | 0.041 | 0.590 | 0.602 | 0.344 |
| BayesC (R) | Bayesian linear | MultiGS-R | 0.357 | 0.370 | 0.050 | 0.507 | 0.616 | 0.090 | 0.582 | 0.601 | 0.226 |
| RFR (R) | Machine learning | MultiGS-R | 0.335 | 0.318 | 0.251 | 0.566 | 0.495 | 0.251 | 0.688 | 0.666 | 0.197 |
| SVR (R) | Machine learning | MultiGS-R | 0.072 | 0.255 | 0.313 | 0.519 | 0.644 | 0.339 | 0.438 | 0.471 | -0.657 |
| RKHS (R) | Kernel-based/Machine learning | MultiGS-R | 0.381 | 0.382 | 0.028 | 0.556 | 0.497 | 0.109 | 0.621 | 0.617 | -0.097 |

| Model | Model type | Tool | Days to maturity (DTM) | | | Oil content (OIL) | | | Plant height (PH) | | |
|---|---|---|---|---|---|---|---|---|---|---|---|
| | | | SNP | HAP | PC | SNP | HAP | PC | SNP | HAP | PC |
| RFC (R) | Machine learning | MultiGS-R | 0.364 | 0.350 | 0.155 | 0.385 | 0.462 | 0.080 | 0.517 | 0.513 | -0.433 |
| SVC (R) | Machine learning | MultiGS-R | 0.265 | 0.363 | 0.272 | 0.491 | 0.674 | 0.120 | 0.133 | 0.182 | -0.590 |
| R_RRBLUP | Linear mixed | MultiGS-P | 0.362 | 0.360 | 0.333 | 0.505 | 0.632 | 0.406 | 0.579 | 0.577 | 0.620 |
| R_GBLUP | Linear mixed | MultiGS-P | 0.411 | 0.410 | 0.410 | 0.530 | 0.604 | 0.528 | 0.086 | 0.370 | 0.082 |
| RRBLUP | Linear mixed | MultiGS-P | 0.318 | 0.333 | 0.257 | 0.507 | 0.630 | 0.338 | 0.557 | 0.558 | 0.563 |
| ElasticNet | Linear | MultiGS-P | 0.304 | 0.258 | 0.291 | 0.672 | 0.561 | 0.450 | 0.679 | 0.651 | 0.631 |
| BRR | Bayesian linear regression | MultiGS-P | 0.361 | 0.359 | 0.333 | 0.561 | 0.633 | 0.406 | 0.578 | 0.576 | 0.620 |
| RFR | Ensemble of trees | MultiGS-P | 0.359 | 0.348 | 0.277 | 0.522 | 0.470 | 0.638 | 0.581 | 0.434 | 0.125 |
| XGBoost | Gradient boosting trees | MultiGS-P | 0.239 | 0.164 | 0.030 | 0.578 | 0.344 | 0.619 | 0.537 | 0.519 | 0.235 |
| LightGBM | Gradient boosting trees | MultiGS-P | 0.217 | 0.181 | 0.001 | 0.559 | 0.260 | 0.491 | 0.657 | 0.629 | 0.442 |
| DNNGS | Deep learning | MultiGS-P | 0.352 | 0.294 | 0.318 | 0.747 | 0.668 | 0.211 | 0.699 | 0.615 | 0.590 |
| HybridAttnMLP | Deep learning | MultiGS-P | 0.256 | 0.329 | 0.019 | 0.445 | 0.605 | 0.400 | 0.536 | 0.632 | 0.562 |
| MLPGS | Deep learning | MultiGS-P | 0.265 | 0.335 | 0.313 | 0.559 | 0.598 | 0.591 | 0.604 | 0.626 | 0.439 |
| GraphConvGS | Deep learning | MultiGS-P | -0.363 | 0.201 | 0.183 | -0.486 | 0.896 | 0.578 | 0.569 | 0.558 | 0.700 |

| Model | Model type | Tool | Days to maturity (DTM) | | | Oil content (OIL) | | | Plant height (PH) | | |
|---|---|---|---|---|---|---|---|---|---|---|---|
| | | | SNP | HAP | PC | SNP | HAP | PC | SNP | HAP | PC |
| GraphAttnGS | Deep learning | MultiGS-P | 0.164 | 0.342 | -0.340 | 0.496 | 0.678 | 0.812 | 0.502 | 0.452 | 0.183 |
| GraphSAGEGS | Deep learning | MultiGS-P | -0.104 | 0.238 | 0.333 | 0.695 | 0.730 | 0.756 | 0.486 | 0.567 | 0.470 |
| GraphFormer | Deep learning | MultiGS-P | 0.212 | 0.299 | 0.363 | 0.655 | 0.691 | 0.698 | 0.661 | 0.564 | 0.574 |
| DeepResBLUP | Deep learning | MultiGS-P | 0.218 | 0.337 | 0.220 | 0.519 | 0.643 | 0.440 | 0.590 | 0.589 | 0.577 |
| DeepBLUP | Deep learning | MultiGS-P | 0.123 | 0.309 | -0.040 | 0.522 | 0.593 | 0.509 | 0.569 | 0.682 | 0.664 |
| EnsembleGS | Deep learning | MultiGS-P | -0.040 | 0.248 | 0.097 | 0.673 | 0.685 | 0.726 | 0.608 | 0.618 | 0.493 |
| DeepGS | Deep learning | Third_party | 0.390 | NA | NA | 0.326 | NA | NA | -0.084 | NA | NA |
| CropFormer | Deep learning | Third_party | -0.068 | NA | NA | 0.001 | NA | NA | 0.401 | NA | NA |
| DPCFormer | Deep learning | Third_party | 0.133 | NA | NA | 0.565 | NA | NA | 0.446 | NA | NA |
| WheatGP | Deep learning | Third_party | 0.000 | NA | NA | 0.034 | NA | NA | -0.147 | NA | NA |
